# Spider mite genotypes with higher growth rate suffer more from competition but exert stronger reproductive interference

**DOI:** 10.1101/2025.08.11.669700

**Authors:** Miguel A. Cruz, Flore Zélé, Raquel Gaspar, Ricardo Santos, Leonor R. Rodrigues, Vitor C. Sousa, Sara Magalhães

**Affiliations:** Centre for Ecology, Evolution and Environmental Changes (CE3C) & CHANGE - Global Change and Sustainability Institute, Faculdade de Ciências da Universidade de Lisboa, Lisboa, Portugal; ISEM, Université de Montpellier, CNRS, IRD, Montpellier, France; Departamento de Biologia Vegetal, Faculdade de Ciências da Universidade de Lisboa, Lisboa, Portugal; Departamento de Biologia Animal, Faculdade de Ciências da Universidade de Lisboa, Lisboa, Portugal; Wissenschaftskolleg zu Berlin, Institute for Advanced Study, Berlin, Germany

**Keywords:** Broad-sense heritability, body size, food competition, genetic correlations, individual variation, reproductive interference

## Abstract

1. Within-species individual variation has been gaining increased attention in ecological studies of species interactions. Yet, these studies rarely consider genetic variation and potential genetic correlations among traits. Moreover, traits beyond those associated with trophic interactions are mostly overlooked. Filling these gaps is important to fully understand the relevance of trait variation in both the ecology and evolution of species interactions.
2. We addressed this using inbred lines of *Tetranychus cinnabarinus*, a spider mite species that often engages in antagonistic interactions with its congeneric *T. urticae*. Specifically, we measured (a) the intrinsic growth rate and the sex ratio in the absence of food competition and reproductive interference; (b) food competition against a density gradient of con- or heterospecific competitors in absence of reproductive interference; (c) reproductive interference in absence of food competition; and (d) the body size of adult females and males.
3. We found significant genetic variance (*i.e.*, broad-sense heritability) among lines for all variables except for the effect of *T. cinnabarinus* on its heterospecific competitor via food competition. Moreover, we showed that lines with higher intrinsic growth were more sensitive to intra- and inter-specific competition for food but exerted stronger reproductive interference on heterospecifics. In addition, the higher the proportion of sons produced by a line, the more resilient it was to reproductive interference but also the lower its growth rate. No genetic correlations were found between any variable and body size.
4. Our results indicate that considering the genetic correlations among traits involved in both trophic and sexual interactions is key to understanding the role of individual variation in species interactions and their evolution.

## Introduction

While community ecology has traditionally treated species as homogenous units based on mean trait values, approaches that acknowledge the importance of within-species variation for ecologically relevant traits have begun to gain traction (Jung et al. 2010; Bolnick et al. 2011; Violle et al. 2012; Des Roches et al. 2018; Westerband et al. 2021). Notably, numerous theoretical studies over the past two decades have investigated the effects of within-species individual variation on community assembly and attempted to integrate them into coexistence theory, often producing conflicting results (reviewed in Stump et al. 2022). Depending on the specific trait being modelled, and whether individual trait variation is present for one or more species, such variation can either promote (*e.g.*, Lichstein et al. 2007; Crawford et al. 2019; Milles et al. 2020), hamper (*e.g.*, Courbaud et al. 2012; Hart et al. 2016; Stump et al. 2020), or have no impact (Banitz 2019) on species coexistence. For example, individual variation in the best competitor is expected to hamper coexistence, whereas it may foster it when present in the worse competitor (Hart et a. 2016, Lichstein et al. 2007). Although attempts have been made to reconcile such disparate results under a broad framework (Stump et al. 2022), accurately assessing the impact of individual variation on species coexistence may require accounting for traits other than those directly related to competition for food, such as traits involved in reproductive interactions.

Negative sexual interactions between species, known as reproductive interference (Gröning and Hochkirch 2008), have garnered recognition as important determinants of species coexistence (*e.g.*, Kishi 2015; Kyogoku 2015, 2020; Grether et al. 2017, 2024; Gómez-Llano et al. 2021). In particular, many empirical studies have shown that reproductive interference can hamper coexistence between ecologically similar species (*e.g.*, Hochkirch et al. 2007; Kishi 2015; Ting and Cutter 2018; Kyogoku et al. 2019; Gómez-Llano et al. 2023). Theoretical work has aimed at integrating these interactions into coexistence theory (*e.g.*, Yoshimura and Clark 1994; Kishi and Nakazawa 2013; Schreiber et al. 2019; Kyogoku and Kokko 2020; Yamamichi et al. 2023), predicting that reproductive interference promotes coexistence under certain conditions, such as when competition for food and reproductive interference are asymmetric and affect each species in opposite ways (Kishi and Nakazawa 2013; Schreiber et al. 2019). In addition, as for traits involved in trophic interactions, individual variation in reproductive interference may also affect coexistence. For example, within-population variation for traits such as mate preferences (*e.g.*, Widemo et al. 1999) and reproductive strategies (Gross 1996) may result in lower levels of reproductive interference for some individuals (by reducing the competition with heterospecifics for certain mate phenotypes or the overlap between reproductively active stages of two species), possibly fostering coexistence.

In addition to the relevance of individual variation for coexistence in general, the role of this variation may be even more impactful when it has a genetic basis. Indeed, some theoretical models predict that, unlike predictions for non-heritable phenotypic variance (Hart et al. 2016), coexistence among competitors is possible when genetic variance for traits in all species is low (Senthilnathan and Gavrilets 2021). Moreover, genetic variance can contribute to species coexistence via its effect on the evolutionary trajectories of populations, for instance through frequency-dependent selection, favouring either one or the other species as they become rare (Vasseur et al. 2011), or by favouring greater resilience to environmental perturbations in the two species (Barabás and D’Andrea 2016).

The impact of heritable trait variation on species coexistence may strongly depend on the strength and direction of potential correlations between traits (Schreiber et al. 2018). Empirical studies have provided evidence of such correlations. For example, Lankau and Strauss (2007) found a negative correlation between intra- and interspecific competition in *Brassica nigra*. Negative correlations have also been found between intrinsic growth and sensitivity to competition in aquatic plants and Bryozoa (Hart et al. 2019; Pettersen et al. 2020). These negative correlations may underlie coexistence between these species and their competitors, as the increasing population sizes of species with higher intrinsic growth can be buffered by their higher sensitivity to competition against heterospecifics (Lankau and Strauss 2007, Hart et al. 2019; Pettersen et al. 2020). Genetic correlations may also promote the evolution of strong competitors, for instance if individuals that are less affected by competition with a heterospecific also exert a stronger competitive pressure on this competitor (Sakarchi and Germain 2023). However, the potential genetic correlations between traits underlying competition for food and reproductive interference remain unaddressed, despite the recognition that such correlations may strongly impact the probability of coexistence between species (Leon 1974; Pease 1984; Lankau 2009, 2011; Weber and Strauss 2016) as well as their evolutionary trajectories (Wilson 2014; Rotter and Holeski 2018).

Here, we set out to test for potential genetic correlations between traits involved in competition for food and in reproductive interference, which can directly bear upon species’ likelihood to coexist. To do so, we assessed broad-sense heritability for such traits and potential correlations among them in a system comprised of two closely related spider mite species, *Tetranychus cinnabarinus* and *T. urticae*. These species are also often referred to as the red and green forms of *T. urticae*, as reproductive isolation is incomplete and gene flow can occur between some populations of the two forms (Auger et al. 2013). However, complete post-zygotic isolation between these forms has also been found, in particular between the populations used here, due to a combination of hybrid sterility and breakdown (Cruz et al. 2021). These two species have overlapping worldwide distributions, share the same host plant species (Migeon and Dorkeld 2023) and often the same individual plant (Lu et al. 2017; Zélé et al. 2018). Moreover, mites are passive dispersers (Clotuche et al. 2013) and may land on plants already colonized by competitors, being thus exposed to contrasting densities of competitors depending on the age of the population. Due to these frequent encounters, they may often engage in competition for food (Lu et al. 2017, 2018) and sexual interactions (Takafuji et al. 1997) in natural populations. This has been confirmed in a laboratory setting, where *T. urticae* excludes *T. cinnabarinus* through a combination of higher population growth when sharing food resources and costly heterospecific matings. Indeed, *T. urticae* males prefer to mate with *T. cinnabarinus* females (Cruz et al. 2025b), resulting in the production of sterile hybrids, which, in combination with first-male sperm precedence (*i.e.*, the first male to mate with a female sires all of its offspring; Helle 1967; Rodrigues et al. 2020), makes this heterospecific cross particularly costly (Cruz et al. 2025b). In contrast, *T. urticae* females that mate with heterospecific males produce male-biased offspring, likely due to gametic incompatibilities leading to fertilization failure (Cruz et al. 2021, 2025a). Indeed, spider mites are haplodiploid, hence unfertilized offspring develop into males. Also, the optimal sex ratio in these species is generally female-biased, hence male production is also costly (Cruz et al. 2021).

Here, we measured food competition and reproductive interference of inbred lines of *T. cinnabarinus* exposed to a single *T. urticae* line, as well as the growth rate, sex ratio and adult body size of *T. cinnabarinus* females and males in absence of heterospecifics. Then, we estimated the broad-sense heritability of these traits (Falconer and Mackay 1996) and tested for correlations among the traits found to be significantly heritable. We discuss the potential implications of such genetic correlations can have on species coexistence and their evolutionary trajectories.

## Methods

### Establishment of spider mite inbred lines

The source population used to establish the focal inbred lines was a genetically diverse population of *T. cinnabarinus* (abbreviated ‘Tc’ hereafter; population ‘*Wu.SS*’, cured from endosymbionts, in Costa et al. 2023) originally created by merging six field-collected populations, as described in Rodrigues et al. (2022). We also used an inbred line of *T. urticae* (‘Tu’), created from a source population established in 2010 in the laboratory (Cruz et al 2021, 2025) previously used to assess the ecological consequences of competition for food and reproductive interference against Tc (Cruz et al. 2025a). Inbred lines were derived from each population following the sib-mating procedure described in Godinho and colleagues (2020), with some adjustments. Briefly, 200 mated adult females of Tc and 20 of Tu were initially isolated on individual *ca*. 9 cm^2^ bean leaf discs pressed onto water-soaked cotton in a Petri dish and allowed to lay eggs for 48h, after which they were discarded. The offspring were then allowed to develop until adulthood on each leaf disc, at which point mating occurred between siblings. From each leaf disc, 14 days after the installation of the initial female, 3 adult sib-mated females were isolated on individual leaf discs to establish the next generation, such that each maternal lineage consisted of 3 replicate discs to reduce the probability of it being lost. The procedure was then repeated as above. At each generation, a single replicate disc was selected per line to establish the next generation. After 15 generations of sib-mating, corresponding to a 93.6% probability of each line being fully inbred (Godinho et al. 2020), we obtained a total of 46 inbred lines of Tc and 9 inbred lines of Tu. Of these, 29 Tc lines and 1 Tu line were randomly selected for this study.

Lines were subsequently maintained on larger bean leaf patches at small population sizes (*ca*. 20 to 70 individuals) under the same controlled laboratory conditions as for the experiments (24±2°C, 70% humidity, 16/8h L/D). This study did not require ethical approval according to the Portuguese decree-law n° 113/2013 and European Union directive 2010/63/EU, which requires this for studies on vertebrates and cephalopods only.

### Measuring traits

We performed two distinct experiments, aimed at quantifying sensitivity to competition and reproductive interference following procedures similar to those detailed in Cruz et al. (2025a). To quantify sensitivity to competition, defined as a reduction in the growth rate (*i.e.*, production of adult daughters) of an individual growing in the presence of others, we measured, without allowing reproductive interference, (a) the growth rate of individuals of each Tc line that were either isolated or exposed to a gradient of intraspecific individuals from the same line or to interspecific individuals from the Tu line, and (b) the growth rate of individuals of the Tu line that were either isolated or exposed to a gradient of intraspecific individuals from the same line or to interspecific individuals from each of the Tc lines. To quantify the sensitivity of Tc, and the harm it caused via reproductive interference with Tu, that is, reductions in growth rate resulting from reproductive interactions with heterospecifics, we measured, without allowing food competition, (a) the growth rate of each Tc line, for individuals that were isolated or exposed to reproductive interference induced by the Tu line and (b) the growth rate of individuals of the Tu line when exposed to reproductive interference by each Tc line. From this second experiment, we also measured the total number of sons, the total number of offspring and the offspring sex ratio. Finally, the body size of females and males of 18 of the Tc lines and of the Tu line was measured as detailed in the Supplementary Materials II.

### Age cohorts

Prior to each experiment, lines were expanded by installing *ca*. 20 adult females of each line in individual small population cages (16 x 16 x 11 cm) containing 4 bean leaves. From these cages, we collected females to form age cohorts in each experiment. For the FC (‘Food competition’) experiment (see the following subsection), we let mated females lay eggs on large leaf fragments placed on water-soaked cotton in separate Petri dishes, such that a sufficient number of mated adult daughters would be available for the experiment 12 days later (*i.e.*, average generation time in standard laboratory conditions, i.e., 24±2°C, 70% humidity, 16/8h L/D). For the RI (‘Reproductive interference’) experiment, this procedure was adapted to obtain either virgin males or virgin females. To obtain males, we isolated 100 (2 * 50) females undergoing their last moulting stage (‘quiescent’ females hereafter) in the absence of males. Those females will remain virgin and thus produce only sons due to spider mite arrhenotoky. Those sons reach adulthood circa 8 days later. To obtain virgin females, we isolated and 150 (3 * 50) adult mated females, which produce both daughters and sons. In both cases, females were isolated on large leaf fragments placed on water-soaked cotton in separate Petri dishes 11 days prior to the experiment. Tu virgin males were directly taken from the age cohorts descending from virgin females on the day of the experiment, whereas quiescent Tu females were isolated two days prior to the experiment), such that they remained virgin until being used as adults in the experiment. As only a small number of individuals (10 females and 10 males) were needed from each Tc line, these were collected directly from their respective small breeding cages.

### Food competition in absence of reproductive interference (‘FC experiment’)

To quantify the sensitivity of Tc lines to food competition with Tu, we placed one focal Tc mated female of each line alone (0 competitors; treatment 1), or with 2, 7 or 19 Tc or Tu competitors (treatments 2 to 7). Conversely, to quantify the harm caused by Tc lines on Tu via food competition, we placed one Tu mated female from the single Tu line alone (treatment 8) or with 2, 7 or 19 Tc individuals from each line (treatments 9 to 11). All treatments were installed on a bean leaf disc with standard size of (9 cm^2^). Fourteen days later, the female offspring in each disc were counted. In this system, competition for food mainly affects developing offspring, causing increased mortality prior to reaching adulthood (Godinho et al. 2023). As all 29 Tc lines could not be tested simultaneously, one experimental replicate of each density of conspecific or heterospecific competitors and of the treatment with the focal female alone were performed for a total of 20 Tc lines and the Tu line in a single temporal block. A total of 15 experimental blocks were carried out, resulting in 10 replicates for each treatment that included the Tc lines and 15 replicates of the treatment where Tu females were tested alone.

### Reproductive interference in absence of food competition (‘RI experiment’)

To quantify the strength of reproductive interference, 5 virgin females of each Tc line and 5 virgin females of the Tu line were given the opportunity to mate with 5 conspecific and 5 heterospecific males for 48 hours on 9 cm^2^ bean leaf discs (‘interaction patches’) before being isolated on 2.5 cm^2^ bean leaf discs (’food patches’) where they laid eggs and their offspring developed. Eight to nine days later, when daughters were still undergoing their last moulting stage, but some offspring sons had already reached adulthood (as spider mite males generally develop faster than females; Mitchell 1973), the offspring from each food patch that originated from the same interaction patch, were merged altogether on a new fresh interaction patch. Two days later 10 now-mated females were randomly isolated on new fresh food patches to lay eggs and for their offspring to develop. Then, the number of adult offspring (sons and daughters) of each isolated female was counted. Hence, growth rate was estimated over two generations to account for the costs resulting from the production of sterile F1 hybrids. This procedure thus allows measuring the impact of reproductive interference while reducing the temporal exposure to competition for food to its minimum. The same procedure was performed for each Tc line and the Tu line in the absence of heterospecifics, such that only 5 females and males of each species were initially installed (single species control treatments). This allowed determining the intrinsic growth rate (daily daughter production per capita) and offspring sex ratio (proportion of sons in the brood) of each line in absence of reproductive interference. The experiment was carried out across 10 independent experimental blocks, with one experimental replicate of each treatment (both species together, Tc alone, and Tu alone) and Tc line per block. Hence, 10 experimental replicates were obtained per treatment per line.

### Theoretical approach for parameter estimation

Interaction coefficients were estimated from data obtained for the different treatments in each of the above experiments, as described in detail in Supplementary Section S4 in Cruz et al. (2025a). Briefly, for the purpose of parameter estimation, we adapted a discrete-time population model (Schreiber et al. 2019), which predicts the number of individuals of species *i* (*N_i_*) that interact through food competition and reproductive interference with individuals of species *j* (*N_j_*), at each generation *t*:

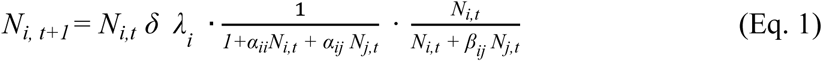

The leftmost term of the expression (*N_i,t_ δ λ_i_*) represents population growth in absence of limiting interactions, where *δ* is the oviposition period in days and *λ_i_* is the per capita daily intrinsic growth rate of species *i*. The middle term 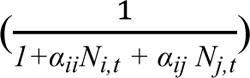 represents the effect of food *1+αiiNi,t + αij Nj,t* competition, where *α_ii_* and *α_ij_* are, respectively, the per capita effects of food competition with *N_i,t_* competitors of species *i* and *N_j,t_* competitors of species *j* (*i.e.*, intraspecific and interspecific competition, respectively) at generation *t*. The rightmost term 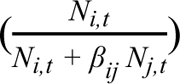 represents the per *Ni,t + β_ij_ Nj,t* capita effect of reproductive interference, where *β_ij_* is the per capita effect of interspecific sexual interactions with *N_j,t_* individuals of species *j* at generation *t*.

The procedure for parameter estimation followed here differs from the one used in Cruz et al. (2025a) in that we used the optimization method ‘L-BFGS-B’, which enables bounding parameter values. Here, for all *α* and *β*, we did not set any maximum value, but a minimum value of 0.0001 to prevent the estimation from converging to negative values (indicative of facilitation; Godoy and Levine 2014), as these are not supported by the framework proposed by Schreiber and colleagues.

### Statistical analyses

Analyses were carried out using the R statistical software (v4.2.2). To assess genetic variance for the effect of trophic and sexual interactions on Tc population growth, we estimated the broad-sense heritability (*H^2^*) of each parameter (*λ_Tc_*, *α_TcTc_*, *α_TcTu_*, *α_TuTc_*, *β_TcTu_*, and *β_TuTc_*) as well as of three other variables measured in the single species treatments of the RI experiment: the number of adult sons produced; the total offspring production, computed as the sum of adult sons and daughters; and the offspring sex ratio, computed as the proportion of sons (*i.e.*, number of sons divided by total offspring production; see Supplementary Table SI1 and Figure SI1, SI2). To obtain *H^2^* estimates, we fitted generalized linear mixed models with a Markov chain Monte Carlo approach using the *MCMCglmm* package (Hadfield 2010) for each combination of focal and competitor individuals (see Table SI2 for detailed information). The daily per capita daughter production (for *λ*, *α* and *β*), number of sons, proportion of sons (*i.e.*, offspring sex ratio), or total adult offspring (sum of sons and daughters), were fit as log-transformed response variables, with a Gaussian error structure, into the models, whereas Tc line identity and experimental block were fit as random explanatory variables. For the FC experiment, where lines were tested under different densities of competitors, the total density of individuals on a patch was also fit as a fixed explanatory variable. *H^2^* were then computed using the variances estimated by the models following the formula:

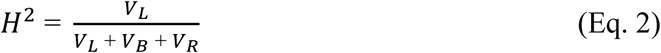

where *V_L_* corresponds to the variance among inbred lines, *V_B_* to the variance among experimental blocks, and *V_R_* to the residual variance of the model. Statistical significance of heritability estimates was determined by comparing the deviance information criterion (DIC) of a model without line identity and another with line identity as a random factor; differences in DIC (ΔDIC) greater than 2 were considered to be significant. All models ran for 300000 iterations, with a burn-in of 10000 iterations and a thinning of 100, and were always provided with a flat, non-informative prior (V=1 and nu=0.002). All models were checked for convergence and auto-correlation, and no issues were found.

Next, we tested for correlations among parameters with significant heritability (*i.e.*, all except *α_TuTc_*; see Figure 1). For each pair of parameters across Tc lines, the observed Pearson correlation coefficient (*r*) was computed with the R function *cor*. Statistical significance was determined by bootstrap permutations (Manly 2018), and the resulting P-values were subjected to an FDR correction to account for multiple testing (Benjamini and Hochberg 1995). For correlations involving the parameter *β_TuTc_*, line 42 was considered an outlier and excluded due to an unusually high mean value and confidence interval.

**Figure 1.**
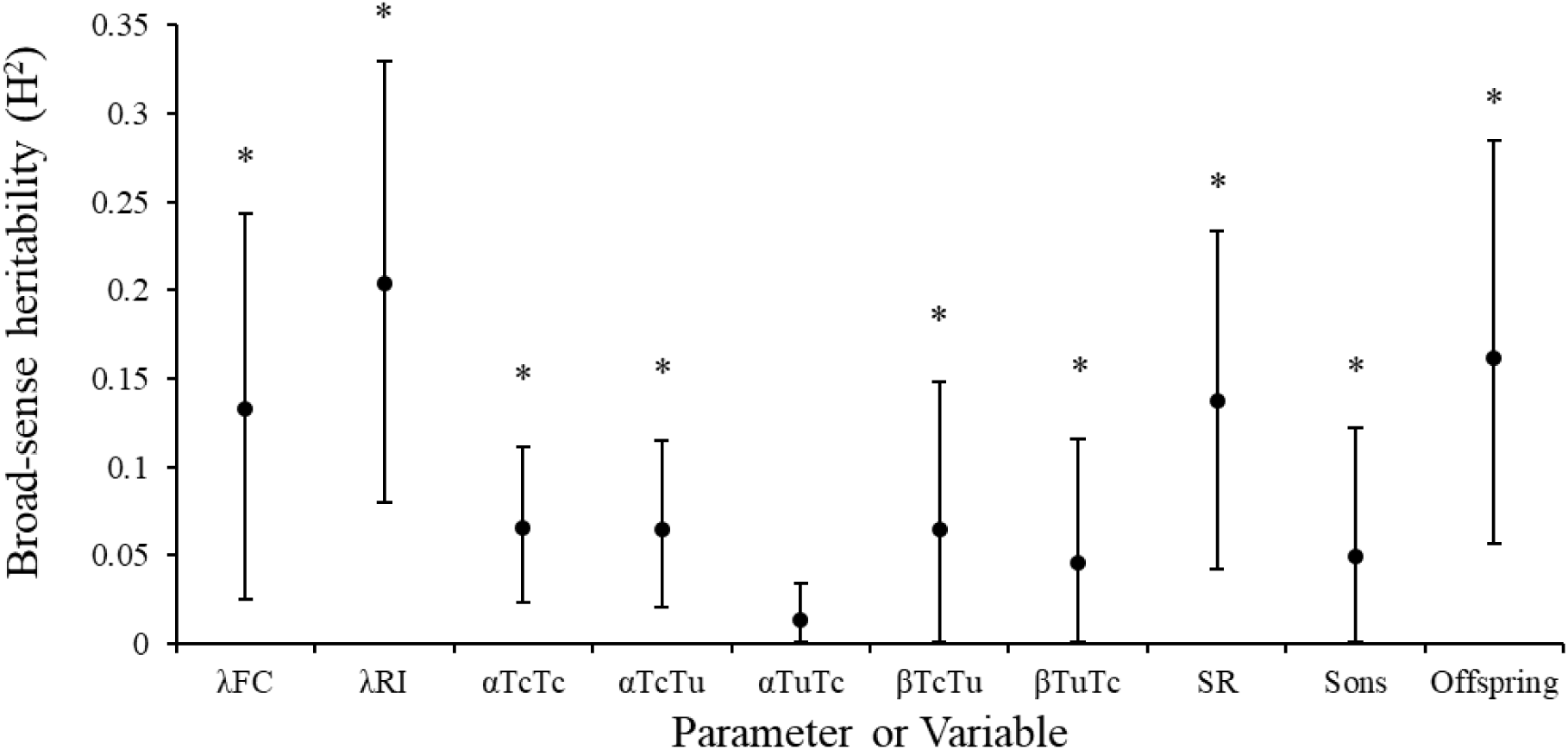
Estimates of heritability for each parameter and variable obtained in the FC and RI experiments. Broad-sense heritability was estimated from the offspring production data obtained for each combination of focal individuals and competitors (FC experiment) or heterospecific mates (RI experiment). Dots represent mean broad-sense heritability for each parameter or variable (see Table SI2 in Supplement I). λ: growth rate; α: sensitivity to competition; β: reproductive interference; Sons: total number of adult sons; Offspring: total number of adult sons and daughters. Error bars represent the 95% highest posterior density interval (HPDI). Asterisks indicate values of heritability significantly different from 0 at the 5% level.

Finally, we assessed *H^2^* of adult female and male body size and tested for genetic correlations with traits involved in trophic and sexual interactions (see Supplementary Material II).

## Results

### Most traits showed significant broad-sense heritability

We found significant broad-sense heritability for the intrinsic growth rate of the Tc lines in both the FC and RI experiments (Figure 1; Table SI2), and for the offspring sex ratio, production of sons, and total production of adult offspring (Figure 1; Table SI2) in the RI experiment. Moreover, significant broad-sense heritability was found for intraspecific and interspecific sensitivity to food competition, and for reproductive interference (Figure 1; Table SI2). There was also significant genetic and/or maternal variance among Tc lines for their negative effect on Tu growth through reproductive interference, but not through competition for food (Figure 1; Table SI2). Finally, female, but not male, body size showed significant broad-sense heritability (see Supplementary Material II).

### Lines with higher intrinsic growth are more sensitive to competition for food

The Tc lines most affected by intraspecific competition (*i.e.*, with individuals of the same line) were also those that were most affected by interspecific competition with the Tu line (*α_TcTc_* and *α_TcTu_*, *r* = 0.86, *P* < 0.0001; Figure 2A; Table SI3). Moreover, both competition coefficients were positively correlated with the intrinsic growth rate of Tc lines (*α_TcTc_* and *λ_FC_*, *r* = 0.89, *P* < 0.0001 and *α_TcTu_* and *λ_FC_*, *r* = 0.78, *P* < 0.0001; Figures 2B, C; Table SI3), indicating a potential trade-off between growth and the ability to cope with competition for food (*i.e.*, the more a line is able to grow in absence of competition, the more it is negatively affected by competitors).

**Figure 2.**
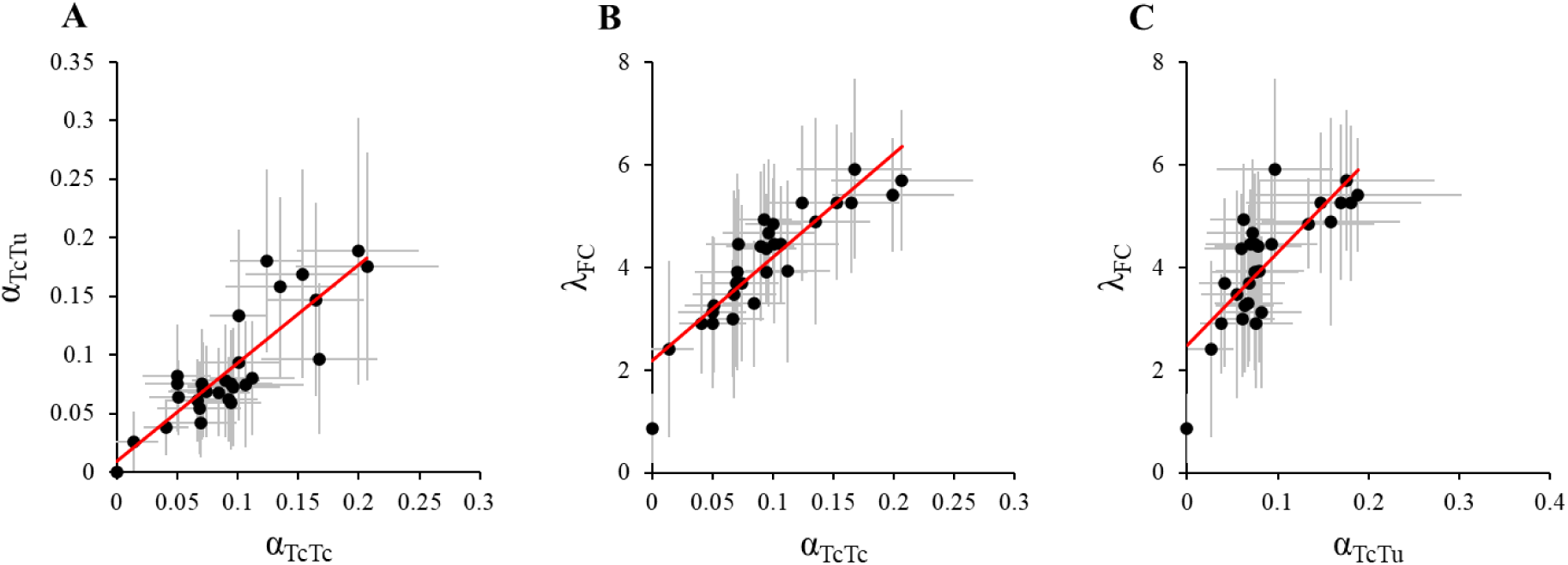
Correlations among parameters estimated from the offspring production data obtained in the FC experiment. Each dot represents, for one Tc line, mean values estimated from the FC experiment for (A) sensitivity to intra- and interspecific competition, α_TcTc_ and α_TcTu_; (B) sensitivity to intraspecific competition, α_TcTc_, and intrinsic growth rates, λ_FC_; and (C) sensitivity to interspecific competition, α_TcTu_, and intrinsic growth rates, λ_FC_. Error bars (in grey) represent the 95% confidence interval. As the sensitivity of Tu to interspecific competition, α_TuTc_, was not heritable, this parameter was not tested for significant correlations. Red trendlines indicate a significant positive correlation between parameters at the 5% level.

### Lines with higher intrinsic growth induce stronger reproductive interference on Tu

The sensitivity to reproductive interference of Tc to Tu was not correlated to that of Tu to Tc (*β_TcTu_* and *β_TuTc_*, *r* = −0.22, *P* = 0.21; Figure 3A; Table SI3). A lack of significant correlation was also found between the sensitivity of Tc lines to reproductive interference and their intrinsic growth rate (*β_TcTu_* and *λ_RI_*, *r* = 0.31, *P* = 0.08; Figure 3B; Table SI3). However, a significant positive correlation was found between the intrinsic growth rate of the Tc lines and the harm they caused to Tu via reproductive interference (*β_TuTc_* and *λ_RI_*, *r* = 0.35, *P* = 0.05; Figure 3B; Table SI3).

**Figure 3.**
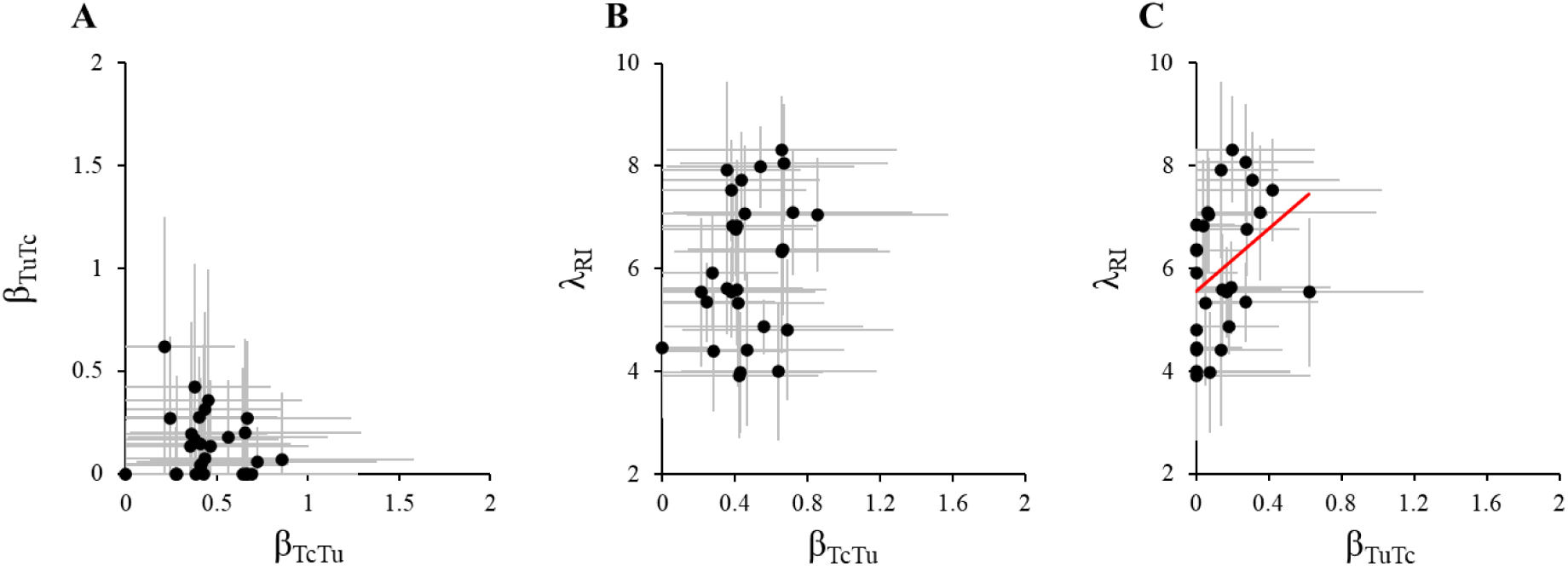
Correlations among parameters estimated from the offspring production data obtained in the RI experiment. Each dot represents, for one Tc line, mean values estimated from the RI experiment for (A) sensitivity to heterospecific mates by Tc, β_TcTu_, and the effect of Tc on heterospecific mates, β_TuTc_; (B) sensitivity to heterospecific mates by Tc, β_TcTu_, and intrinsic growth rates, λ_RI_; and (C) the effect of Tc on heterospecific mates, β_TuTc_, and intrinsic growth rates, λ_RI_. Error bars represent the 95% confidence interval. The red trendline indicates a significant positive correlation between parameters at the 5% level.

### The Tc lines most sensitive to intraspecific competition induce the strongest reproductive interference on Tu

The sensitivity of Tc to reproductive interference by Tu did not significantly correlate with its sensitivity to intraspecific or interspecific competition for food (*β_TcTu_* and *α_TcTc_*, *r* = −0.09, *P* = 0.38 and *β_TcTu_* and *α_TcTu_*, *r* = −0.04, *P* = 0.41, respectively; Figure 4A, B; Table SI3), nor did the latter correlate with the effect of Tc on Tu through reproductive interference (*α_TcTu_* and *β_TuTc_*, *r* = 0.17, *P* = 0.26; Figure 4D; Table SI3). However, the Tc lines most affected by intraspecific competition were also the ones with the greatest negative effect on the Tu line through reproductive interference (*α_TcTc_* and *β_TuTc_*, *r* = 0.42, *P* = 0.04; Figure 4C; Table SI3).

**Figure 4.**
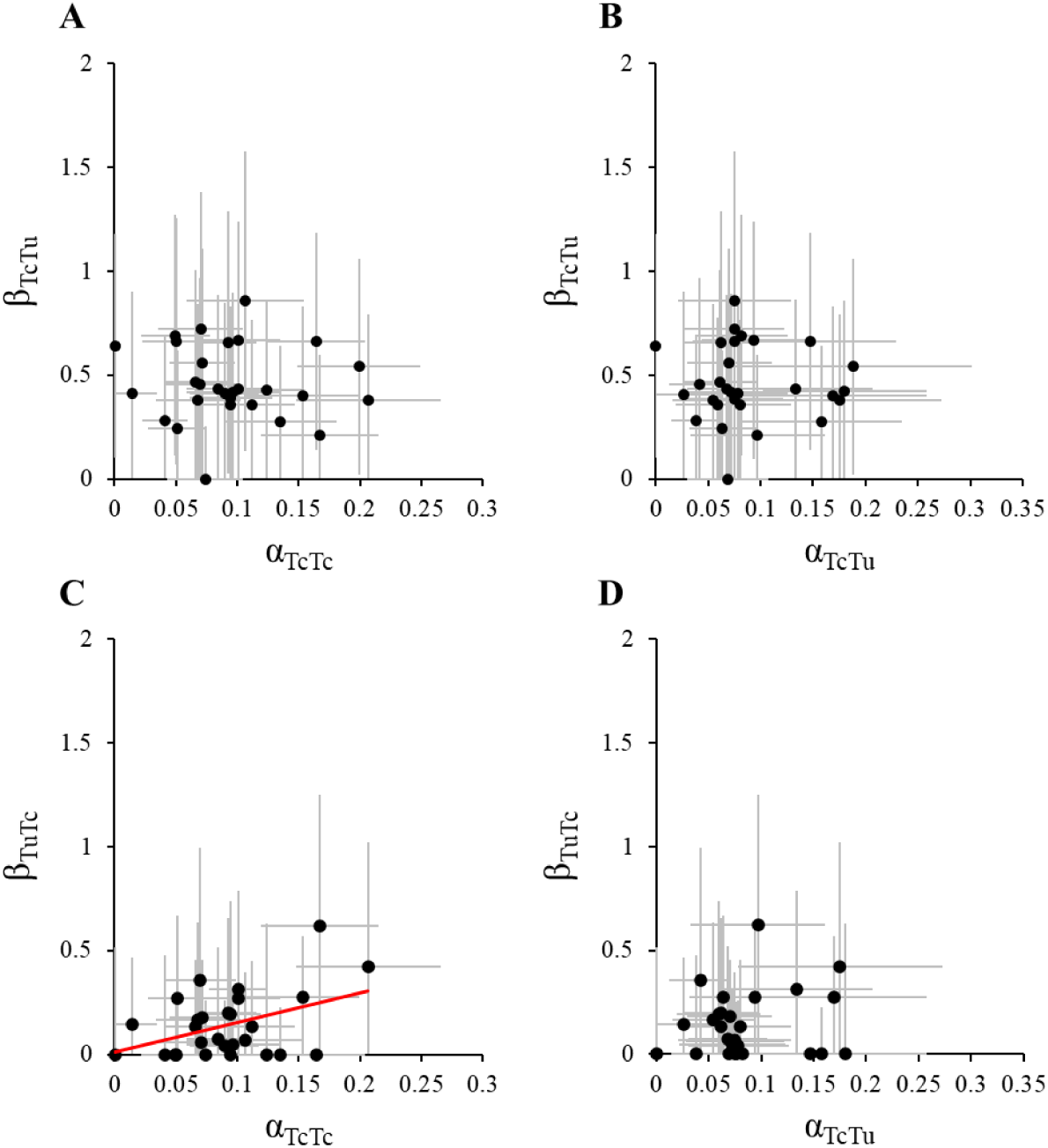
Correlations among parameters related to competition for food and to reproductive interference. Each dot represents, for one Tc line, mean values estimated for (A) sensitivity to conspecific competitors, α_TcTc_, and to heterospecific mates, β_TcTu_; (B) sensitivity to heterospecific competitors, α_TcTu_, and to heterospecific mates, β_TcTu_; (C) sensitivity to conspecific competitors, α_TcTc_, and the effect of Tc on heterospecific mates, β_TuTc_; and (D) sensitivity to heterospecific competitors, α_TcTu_, and the effect of Tc on heterospecifics mates, β_TcTu_. Error bars represent the 95% confidence interval. As the sensitivity of Tu to interspecific competition, α_TuTc_, was found to not be heritable, this parameter was not tested for significant correlations. Red trendlines indicate a significant positive correlation between parameters at the 5% level.

### Correlations between food competition and reproductive interference may be explained by the offspring sex ratio

To investigate a possible explanation for the relationship between reproductive interference and food competition, we tested for correlations between these parameters and the offspring sex ratio. Although the number of daughters produced did not affect the strength of reproductive interference experienced by Tc (see *β_TcTu_* and *λ_RI_*; Figure 3B), we found a significant negative correlation between offspring sex ratio in absence of heterospecifics and the sensitivity of Tc to reproductive interference by Tu (*sex ratio* and *β_TcTu_*, *r* = −0.38, *P* = 0.03; Figure 5A; Table SI3). Thus, independently of the absolute number of females produced, Tc lines that produce a higher proportion of sons were less affected by reproductive interference. This was not due to an increase in the absolute number of sons, as both intrinsic growth and offspring sex ratio correlate with total offspring production (*total offspring* and *λ_RI_*, *r* = 0.96, *P* < 0.0001; and *sex ratio* and *total offspring*, *r* = −0.42, *P* = 0.03, respectively; Figure S4A, B; Table SI3), but total offspring was unrelated to the overall number of sons (*total offspring* and *sons*, *r* = 0.15, *P* = 0.29; Figure S4C; Table SI3).

**Figure 5.**
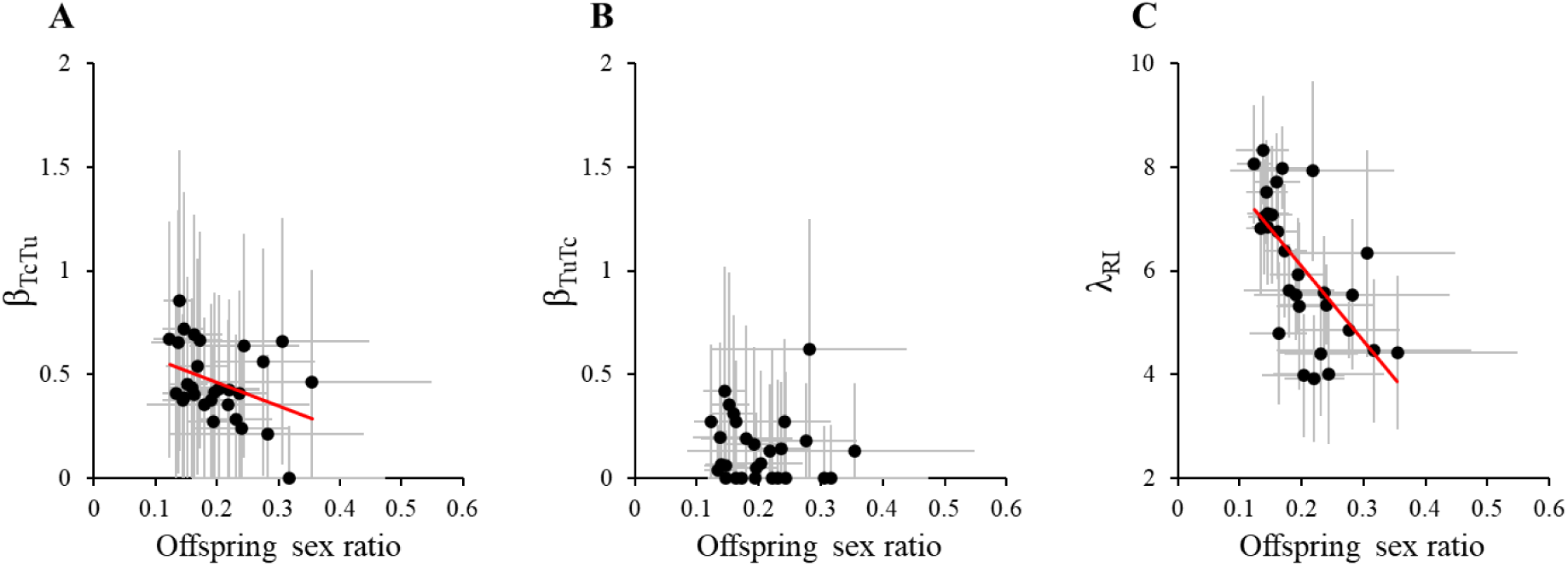
Correlations between offspring sex ratio, sensitivity to, and harm caused via reproductive interference in the RI experiment. Each dot represents, for one Tc line, the mean estimated values of the offspring sex ratio (computed as the proportion of sons) and (A) the sensitivity of Tc to reproductive interference, β_TcTu_, (B) the effect of Tc on Tu via reproductive interference, β_TuTc_, and (C) the intrinsic growth rate of Tc. Error bars represent the 95% confidence interval. The red trendline indicates a significant negative correlation between parameters at the 5% level.

Although we found a positive correlation between the intrinsic growth of the Tc lines and the harm they inflicted on Tu through reproductive interference (see *β_TuTc_* and *λ_RI_*; Figure 3C), offspring sex ratio (the proportion of males) was negatively correlated with the former (*sex ratio* and *λ_RI_*, *r* = −0.64, *P* < 0.0001; Figure 5C; Table SI3), and not correlated with the latter (*sex ratio* and *β_TuTc_*, *r* = −0.07, *P* = 0.39; Figure 5B; Table SI3). Thus, offspring sex ratio did not directly explain the harm caused by Tc to Tu via reproductive interference, but only indirectly through its relationship to the number of daughters produced. Finally, female body size did not correlate with any other variable (see Supplementary Material II).

## Discussion

In this study, we assessed the genetic variance for and potential genetic correlations between interaction coefficients and traits associated with food competition and reproductive interference in inbred lines of the spider mite *T. cinnabarinus* interacting with a line of a closely-related species, *T. urticae*. We found significant genetic (and/or maternal) variance for all estimated parameters except for the sensitivity of Tu to food competition with Tc. Moreover, the sensitivity of Tc lines to intraspecific and to interspecific competition were positively correlated, as were both these sensitivities with the intrinsic growth rate of Tc. Furthermore, all aforementioned parameters were positively correlated with the strength of reproductive interference of Tc on Tu. Finally, the sensitivity of Tc lines to reproductive interference was also found to correlate negatively with the proportion of males they produced in their offspring.

### Genetic variance for biotic interactions was significant, but low

The values of broad-sense heritability estimated for parameters and variables were overall low. This is consistent with the relatively low contributions of genetic (and/or maternal) factors to determining competitive interactions identified in other systems (*e.g.*, Johnson et al. 2008; Tian et al. 2022, but see Annicchiarico et al. 1999; Shuster et al. 2006). These low values could be due to a high environmental variance for these traits, as found for behavioural traits (Dochtermann et al. 2019). Alternatively, these traits may be closely related to fitness, in which case their genetic variance is expected to be low due to the effect of natural selection, which eliminates non-fit genotypes (Hoffmann et al. 2016). The remaining genetic variance that we detected may have been maintained through the existence of trade-offs between traits as predicted by classical quantitative genetics (Falconer and Mackay 1996).

### Intrinsic growth rates correlate with sensitivity to both biotic interactions

The lines that had higher intrinsic growth rates also experienced stronger negative effects of competition for food. This is in agreement with past studies showing trade-offs between intrinsic growth rates and the ability to cope with competition in other systems (*e.g.*, Luckinbill 1979; Hart et al. 2019; Pettersen et al. 2020, but see Sakarchi and Germain 2023). Note that this negative correlation could have been masked if the lines had shown low growth values in absence of competitors. Moreover, this correlation could be due to a negative correlation between body size and number of the offspring, as smaller individuals may suffer more from competition, whereas a larger brood corresponds to a larger growth rate (*e.g.*, Blanckenhorn et al. 2011). However, we found no significant correlation between female or male body size and any of the traits measured, and thus could not provide support for this hypothesis.

We also found a positive correlation between the intrinsic growth rate and the strength of reproductive interference that Tc inflicts on Tu. This may be due to reproductive interference being density dependent, as its strength increases with more encounters with heterospecifics (Kuno 1992; Gröning and Hochkirch 2008; Kyogoku and Sota 2017). However, the lack of a negative correlation between intrinsic growth and sensitivity of Tc to reproductive interference by Tu runs counter to these same expectations. Again, this incongruence may stem from other traits involved in these reproductive interactions, such as the sex ratio.

### Offspring sex ratio as a mediator of correlations among food competition and reproductive interference

Given the important role of sex ratio in both competitive and reproductive interactions, we investigated whether this variable could provide a (partial) explanation for the correlations observed. The offspring sex ratio was negatively correlated with intrinsic growth rate, as Tc lines that produced more offspring also produced fewer sons. Furthermore, lines with a lower proportion of males suffered more from reproductive interference. This suggests that the proportion of males has a direct impact on reproductive interference in this system. Possibly, this is because males guard immature females in order to secure mating opportunities (Satoh et al. 2001), and prefer virgin females over mated ones (Rodrigues et al. 2017), whereas females tend to show no mating preferences (e.g., Rodrigues et al. 2022). Given first-male sperm precedence in this species (Rodrigues et al. 2020), a first mating with a conspecific male is expected to protect females from the negative impact of reproductive interference in the long-term. Therefore, in spider mites, the relative proportion of conspecific to heterospecific males is expected to be an important factor in determining vulnerability to heterospecific matings (as in frogs; Hettyey and Pearman 2003).

### The effect of genetic correlations and individual variation on coexistence

Coexistence between species competing for shared resources is expected to be facilitated by a trade-off between intra- and interspecific competition (Vasseur et al. 2011). Such trade-offs have been shown for both plant and animal species (*e.g.*, Lankau 2008; Liendo et al. 2018), although this is not universal (Miller and Schemske 1990). We did not find this here. Thus, according to theoretical expectations, local coexistence between these mite species is not possible. However, we tested competition between these species in an environment containing only one type of resource and very limited space. Perhaps under conditions allowing for access to more varied resource types and for spatial segregation, which may reduce encounter rates and interactions between the two species, stabilizing mechanisms could arise and foster coexistence even in the absence of a genetic trade-off between intra- and interspecific competition. This is supported by theoretical work which suggests that reproductive interactions between species can drive aggregation and habitat specialization (Nishida et al. 2015; Ruokolainen and Hanski 2016; Kyogoku and Kokko 2020).

Theory also predicts that coexistence between species engaging in both trophic (including competition, an indirect trophic interaction) and sexual interactions should be possible if there is a trade-off between interaction strengths, such that the superior competitor for food is inferior in reproductive interference and vice-versa (Kishi and Nakazawa 2013). Here, we find no evidence of such a trade-off but instead that lines that suffered less from intra-and interspecific food competition inflicted less harm through reproductive interference. Therefore, the genetic correlations we detected are expected to hamper, rather than to foster, coexistence. However, we also show that competition traits in Tc, the inferior competitor in this system (Cruz et al 2025a), exhibit variation. This is expected to instead promote coexistence, except if variation in the superior competitor is also large (Begon and Wall 1987; Lichstein et al. 2007; Hart et al. 2016), a hypothesis that would require investigating variation for these traits in Tu as well. Thus, our findings set the stage for future experimental studies investigating the impact of individual variation on coexistence between species engaging in both food competition and reproductive interference.

In sum, this study shows that traits associated to competition for food and reproductive interference are not independent, which should be considered in future studies investigating species distributions and the evolutionary trajectories of their populations.

## Acknowledgements

We are grateful to Juan Lopez and Arianna Thomas-Cabianca for assistance with the maintenance of cultures and inbred lines and in data collection, and to Inês Fragata, João Picão Osório and Sofia Henriques for advice regarding the statistical analyses. This work was funded by an ERC Consolidator Grant (COMPCON, GA 725419) attributed to SM and by Fundação para a Ciência e Tecnologia (FCT) through CE3C - Centre for Ecology, Evolution and Environmental Changes unit funding (UIDB/00329/2025). MAC, VCS and SM acknowledge financial support provided by the FCT project GeneticBarriers (2022.03475.PTDC; PI: Alexandre Blanckaert). MAC was funded through an FCT PhD fellowship (SFRH/BD/136454/2018) and VCS and LRR by FCT contracts (CEECINST/00032/2018/CP1523/CT0008 and <u>CEECIND/CP1715/CT0007</u>). This is contribution ISEM-XXX of the Institute of Evolutionary Science of Montpellier (ISEM). Funding agencies did not participate in the design or analysis of experiments. For the purpose of Open Access, a CC-BY 4.0 public copyright licence has been applied by the authors to the present document and will be applied to all subsequent versions up to the Author Accepted Manuscript arising from this submission.

## References

Annicchiarico, P., E. Piano, and I. Rhodes. 1999. Heritability of, and genetic correlations among, forage and seed yield traits in Ladino white clover. Plant Breed. 118:341–346.

Auger, P., A. Migeon, E. A. Ueckermann, L. Tiedt, and M. Navajas. 2013. Evidence for synonymy between *Tetranychus urticae* and *Tetranychus cinnabarinus* (Acari, Prostigmata, Tetranychidae): review and new data. Acarologia 53:383–415.

Banitz, T. 2019. Spatially structured intraspecific trait variation can foster biodiversity in disturbed, heterogeneous environments. Oikos 128:1478–1491.

Barabás, G., and R. D’Andrea. 2016. The effect of intraspecific variation and heritability on community pattern and robustness. Ecol. Lett. 19:977–986.

Begon, M., and R. Wall. 1987. Individual Variation and Competitor Coexistence: A Model. Funct. Ecol. 1:237–241.

Benjamini, Y., and Y. Hochberg. 1995. Controlling the False Discovery Rate: A Practical and Powerful Approach to Multiple Testing. J. R. Stat. Soc. Ser. B Stat. Methodol. 57:289–300.

Blanckenhorn, W. U., P. E. A. Hoeck, C. Reim, and Y. Teuschl. 2011. A cost of being large: genetically large yellow dung flies lose out in intra-specific food competition. Evol. Ecol. 25:875–884.

Bolnick, D. I., P. Amarasekare, M. S. Araújo, R. Bürger, J. M. Levine, M. Novak, V. H. W. Rudolf, S. J. Schreiber, M. C. Urban, and D. A. Vasseur. 2011. Why intraspecific trait variation matters in community ecology. Trends Ecol. Evol. 26:183–192.

Christie, K., and S. Y. Strauss. 2020. Frequency-dependent fitness and reproductive dynamics contribute to habitat segregation in sympatric jewelflowers. Proc. R. Soc. B Biol. Sci. 287:20200559.

Clotuche, G., M. Navajas, A.-C. Mailleux, and T. Hance. 2013. Reaching the Ball or Missing the Flight? Collective Dispersal in the Two-Spotted Spider Mite Tetranychus urticae. PLOS ONE 8:e77573.

Costa, S. G., S. Magalhães, and L. R. Rodrigues. 2023. Multiple mating rescues offspring sex ratio but not productivity in a haplodiploid exposed to developmental heat stress. Funct. Ecol. 37:1291–1303.

Courbaud, B., G. Vieilledent, and G. Kunstler. 2012. Intra-specific variability and the competition–colonisation trade-off: coexistence, abundance and stability patterns. Theor. Ecol. 5:61–71.

Crawford, M., F. Jeltsch, F. May, V. Grimm, and U. E. Schlägel. 2019. Intraspecific trait variation increases species diversity in a trait-based grassland model. Oikos 128:441–455.

Cruz, M. A., O. Godoy, I. Fragata, V. C. Sousa, S. Magalhães, and F. Zélé. 2025a. Competition for food affects the strength of reproductive interference and its consequences for species coexistence. Funct. Ecol. 39:1982–1997.

Cruz, M. A., S. Magalhães, M. Bakırdöven, and F. Zélé. 2025b. *Wolbachia* strengthens the match between premating and early postmating isolation in spider mites. Evolution 79:203–219.

Cruz, M. A., S. Magalhães, É. Sucena, and F. Zélé. 2021. *Wolbachia* and host intrinsic reproductive barriers contribute additively to postmating isolation in spider mites. Evolution 75:2085–2101.

Des Roches, S., D. M. Post, N. E. Turley, J. K. Bailey, A. P. Hendry, M. T. Kinnison, J. A. Schweitzer, and E. P. Palkovacs. 2018. The ecological importance of intraspecific variation. Nat. Ecol. Evol. 2:57–64.

Dochtermann, N. A., T. Schwab, M. Anderson Berdal, J. Dalos, and R. Royauté. 2019. The Heritability of Behavior: A Meta-analysis. J. Hered. 110:403–410.

Falconer, D. S., and T. F. C. Mackay. 1996. Introduction to Quantitative Genetics. 4th ed. Longmans Green, Harlow, Essex, UK.

Godinho, D. P., M. A. Cruz, M. Charlery de la Masselière, J. Teodoro-Paulo, C. Eira, I. Fragata, L. R. Rodrigues, F. Zélé, and S. Magalhães. 2020. Creating outbred and inbred populations in haplodiploids to measure adaptive responses in the laboratory. Ecol. Evol. 10:7291–7305.

Godinho, D. P., L. R. Rodrigues, S. Lefèvre, L. Delteil, A. F. Mira, I. R. Fragata, S. Magalhães, and A. B. Duncan. 2023. Limited host availability disrupts the genetic correlation between virulence and transmission. Evol. Lett. 7:58–66.

Godoy, O., and J. M. Levine. 2014. Phenology effects on invasion success: insights from coupling field experiments to coexistence theory. Ecology 95:726–736.

Gómez-Llano, M., W. A. Boys, T. Ping, S. P. Tye, and A. M. Siepielski. 2023. Interactions between fitness components across the life cycle constrain competitor coexistence. J. Anim. Ecol. 92:2297–2308.

Gómez-Llano, M., R. M. Germain, D. Kyogoku, M. A. McPeek, and A. M. Siepielski. 2021. When ecology fails: how reproductive interactions promote species coexistence. Trends Ecol. Evol. 36:610–622.

Grether, G. F., A. E. Finneran, and J. P. Drury. 2024. Niche differentiation, reproductive interference, and range expansion. Ecol. Lett. 27:e14350.

Grether, G. F., K. S. Peiman, J. A. Tobias, and B. W. Robinson. 2017. Causes and consequences of behavioral interference between species. Trends Ecol. Evol. 32:760–772.

Gröning, J., and A. Hochkirch. 2008. Reproductive interference between animal species. Q. Rev. Biol. 83:257–282.

Gross, M. R. 1996. Alternative reproductive strategies and tactics: diversity within sexes. Trends Ecol. Evol. 11:92–98.

Hadfield, J. D. 2010. MCMC methods for multi-response generalized linear mixed models: The MCMCglmm R package. J. Stat. Softw. 33:1–22.

Hart, S. P., S. J. Schreiber, and J. M. Levine. 2016. How variation between individuals affects species coexistence. Ecol. Lett. 19:825–838.

Hart, S. P., M. M. Turcotte, and J. M. Levine. 2019. Effects of rapid evolution on species coexistence. Proc. Natl. Acad. Sci. 116:2112–2117.

Helle, W. 1967. Fertilization in the Two-Spotted Spider Mite (*Tetranychus urticae*: Acari). Entomol. Exp. Appl. 10:103–110.

Hettyey, A., and P. B. Pearman. 2003. Social environment and reproductive interference affect reproductive success in the frog Rana latastei. Behav. Ecol. 14:294–300.

Hochkirch, A., J. Gröning, and A. Bücker. 2007. Sympatry with the devil: reproductive interference could hamper species coexistence. J. Anim. Ecol. 76:633–642.

Hoffmann, A. A., J. Merilä, and T. N. Kristensen. 2016. Heritability and evolvability of fitness and nonfitness traits: Lessons from livestock. Evolution 70:1770–1779.

Johnson, M. T. J., R. Dinnage, A. Y. Zhou, and M. D. Hunter. 2008. Environmental Variation Has Stronger Effects than Plant Genotype on Competition among Plant Species. J. Ecol. 96:947–955.

Jung, V., C. Violle, C. Mondy, L. Hoffmann, and S. Muller. 2010. Intraspecific variability and trait-based community assembly. J. Ecol. 98:1134–1140.

Kishi, S. 2015. Reproductive interference in laboratory experiments of interspecific competition. Popul. Ecol. 57:283–292.

Kishi, S., and T. Nakazawa. 2013. Analysis of species coexistence co-mediated by resource competition and reproductive interference. Popul. Ecol. 55:305–313.

Kuno, E. 1992. Competitive exclusion through reproductive interference. Popul. Ecol. 34:275–284.

Kyogoku, D. 2015. Reproductive interference: ecological and evolutionary consequences of interspecific promiscuity. Popul. Ecol. 57:253–260.

Kyogoku, D. 2020. When does reproductive interference occur? Predictions and data. Popul. Ecol. 62:196–206.

Kyogoku, D., and H. Kokko. 2020. Species coexist more easily if reinforcement is based on habitat preferences than on species recognition. J. Anim. Ecol. 89:2605–2616.

Kyogoku, D., M. Kondoh, and T. Sota. 2019. Does past evolutionary history under different mating regimes influence the demographic dynamics of interspecific competition? Ecol. Evol. ece3.5397.

Kyogoku, D., and T. Sota. 2017. A generalized population dynamics model for reproductive interference with absolute density dependence. Sci. Rep. 7:1–8.

Lankau, R. 2008. A chemical trait creates a genetic trade-off between intra- and interspecific competitive ability. Ecology 89:1181–1187.

Lankau, R. A. 2009. Genetic Variation Promotes Long-Term Coexistence of Brassica nigra and Its Competitors. Am. Nat. 174:E40–E53.

Lankau, R. A. 2011. Rapid evolutionary change and the coexistence of species. Annu. Rev. Ecol. Evol. Syst. 42:335–354.

Lankau, R. A., and S. Y. Strauss. 2007. Mutual feedbacks maintain both genetic and species diversity in a plant community. Science 317:1561–1563.

Leon, J. A. 1974. Selection in Contexts of Interspecific Competition. Am. Nat. 108:739–757.

Lichstein, J. W., J. Dushoff, S. A. Levin, and S. W. Pacala. 2007. Intraspecific Variation and Species Coexistence. Am. Nat. 170:807–818.

Liendo, M. C., M. A. Parreño, J. L. Cladera, M. T. Vera, and D. F. Segura. 2018. Coexistence between two fruit fly species is supported by the different strength of intra- and interspecific competition. Ecol. Entomol. 43:294–303.

Lu, W., Y. Hu, P. Wei, Q. Xu, C. Bowman, M. Li, and L. He. 2018. Acaricide-mediated competition between the sibling species *Tetranychus cinnabarinus* and *Tetranychus urticae*. J. Econ. Entomol. 111:1346–1353.

Lu, W., M. Wang, Z. Xu, G. Shen, P. Wei, M. Li, W. Reid, and L. He. 2017. Adaptation of acaricide stress facilitates *Tetranychus urticae* expanding against *Tetranychus cinnabarinus* in China. Ecol. Evol. 7:1233–1249.

Luckinbill, L. S. 1979. Selection and the r/K Continuum in Experimental Populations of Protozoa. Am. Nat. 113:427–437.

Manly, B. F. J. 2018. Randomization, Bootstrap and Monte Carlo Methods in Biology. 3rd ed. Chapman and Hall/CRC, New York.

Migeon, A., and F. Dorkeld. 2023.Spider Mites Web: a comprehensive database for the Tetranychidae.

Miller, T. E., and D. W. Schemske. 1990. An Experimental Study of Competitive Performance in *Brassica rapa* (cruciferae). Am. J. Bot. 77:993–998.

Milles, A., M. Dammhahn, and V. Grimm. 2020. Intraspecific trait variation in personality-related movement behavior promotes coexistence. Oikos 129:1441–1454.

Mitchell, R. 1973. Growth and population dynamics of a spider mite (*Tetranychus urticae* K., Acarina: Tetranychidae). Ecology 54:1349–1355.

Nishida, T., K. Takakura, and K. Iwao. 2015. Host specialization by reproductive interference between closely related herbivorous insects. Popul. Ecol. 57:273–281.

Pease, C. M. 1984. On the Evolutionary Reversal of Competitive Dominance. Evolution 38:1099–1115.

Pettersen, A. K., M. D. Hall, C. R. White, and D. J. Marshall. 2020. Metabolic rate, context-dependent selection, and the competition-colonization trade-off. Evol. Lett. 4:333–344.

Rodrigues, L. R., A. R. T. Figueiredo, T. Van Leeuwen, I. Olivieri, and S. Magalhães. 2020. Costs and benefits of multiple mating in a species with first-male sperm precedence. J. Anim. Ecol. 89:1045–1054.

Rodrigues, L. R., A. R. T. Figueiredo, S. A. M. Varela, I. Olivieri, and S. Magalhães. 2017. Male spider mites use chemical cues, but not the female mating interval, to choose between mates. Exp. Appl. Acarol. 71:1–13.

Rodrigues, L. R., F. Zélé, I. Santos, and S. Magalhães. 2022. No evidence for the evolution of mating behavior in spider mites due to *Wolbachia*-induced cytoplasmic incompatibility. Evolution 76:623–635.

Rotter, M. C., and L. M. Holeski. 2018. A meta-analysis of the evolution of increased competitive ability hypothesis: genetic-based trait variation and herbivory resistance trade-offs. Biol. Invasions 20:2647–2660.

Ruokolainen, L., and I. Hanski. 2016. Stable coexistence of ecologically identical species: conspecific aggregation via reproductive interference. J. Anim. Ecol. 85:638–647.

Sakarchi, J., and R. M. Germain. 2023. The evolution of competitive ability. Am. Nat. 201:1–15.

Satoh, Y., S. Yano, and A. Takafuji. 2001. Mating strategy of spider mite, Tetranychus urticae (Acari: Tetranychidae) males: Postcopulatory guarding to assure paternity. Appl. Entomol. Zool. 36:41–45.

Schreiber, S. J., S. Patel, and C. terHorst. 2018. Evolution as a Coexistence Mechanism: Does Genetic Architecture Matter? Am. Nat. 191:407–420.

Schreiber, S. J., M. Yamamichi, and S. Y. Strauss. 2019. When rarity has costs: coexistence under positive frequency-dependence and environmental stochasticity. Ecology 100:1–13.

Senthilnathan, A., and S. Gavrilets. 2021. Ecological consequences of intraspecific variation in coevolutionary systems. Am. Nat. 197:1–17.

Shuster, S. M., E. V. Lonsdorf, G. M. Wimp, J. K. Bailey, and T. G. Whitham. 2006. Community heritability measures the evolutionary consequences of indirect genetic effects on community structure. Evolution 60:991–1003.

Stump, S. M., J. H. Marden, N. G. Beckman, S. A. Mangan, and L. S. Comita. 2020. Resistance Genes Affect How Pathogens Maintain Plant Abundance and Diversity. Am. Nat. 196:472–486.

Stump, S. M., C. Song, S. Saavedra, J. M. Levine, and D. A. Vasseur. 2022. Synthesizing the effects of individual-level variation on coexistence. Ecol. Monogr. 92:e01493.

Takafuji, A., E. Kuno & H. Fujimoto. 1997. Reproductive interference and its consequences for the competitive interactions between two closely related *Panonychus* spider mites. Exp. Appl. Acarol. 21, 379–39.

Tian, X., H. Ohtsuki, and J. Urabe. 2022. Competitive consequences determined by phenotypic but not genetic distance: A study with asexual water flea genotypes. Funct. Ecol. 36:2152–2162.

Ting, J. J., and A. D. Cutter. 2018. Demographic consequences of reproductive interference in multi-species communities. BMC Ecol. 18:1–13.

Vasseur, D. A., P. Amarasekare, V. H. W. Rudolf, and J. M. Levine. 2011. Eco-evolutionary dynamics enable coexistence via neighbor-dependent selection. Am. Nat. 178:E96–E109.

Violle, C., B. J. Enquist, B. J. McGill, L. Jiang, C. H. Albert, C. Hulshof, V. Jung, and J. Messier. 2012. The return of the variance: Intraspecific variability in community ecology. Trends Ecol. Evol. 27:244–252.

Weber, M. G., and S. Y. Strauss. 2016. Coexistence in close relatives: beyond competition and reproductive isolation in sister taxa. Annu. Rev. Ecol. Evol. Syst. 47:359–381.

Westerband, A. C., J. L. Funk, and K. E. Barton. 2021. Intraspecific trait variation in plants: a renewed focus on its role in ecological processes. Ann. Bot. 127:397–410.

Widemo, F., S. A. Sæther, F. Widemo, S. A. Sæther, F. Widemo, S. A. Sæther, F. Widemo, S. A. Sæther, F. Widemo, and S. A. Sæther. 1999. Beauty is in the eye of the beholder: causes and consequences of variation in mating preferences. Trends Ecol. Evol. 14:26–31.

Wilson, A. J. 2014. Competition as a source of constraint on life history evolution in natural populations. Heredity 112:70–78.

Yamamichi, M., K. Tsuji, S. Sakai, and E. I. Svensson. 2023. Frequency-dependent community dynamics driven by sexual interactions. Popul. Ecol. 65:204–219.

Yoshimura, J., and C. W. Clark. 1994. Population dynamics of sexual and resource competition. Theor. Popul. Biol. 45:121–131.

Zélé, F., J. L. Santos, D. P. Godinho, and S. Magalhães. 2018. *Wolbachia* both aids and hampers the performance of spider mites on different host plants. FEMS Microbiol. Ecol. 94:fiy187.

