## Supplementary materials I for "Spider mite genotypes with higher growth rate suffer more from competition but exert stronger reproductive interference"

**Table SI1. Estimated parameter values and offspring production variables obtained in the FC and the RI experiments.** The mean (± 95% confidence interval) parameter values estimated from the models fit to offspring data of the FC and the RI experiments are provided for each Tc line. The intrinsic growth rate was estimated from both experiments (hence $\lambda_{FC}$ and $\lambda_{RI}$); the competition parameters $\alpha_{TcTc}$, $\alpha_{TuTc}$ and $\alpha_{TcTu}$ (which describe, respectively, the competitive effect that each Tc line has on itself, on the Tu line, and that the Tu line has on each Tc line) were estimated from the FC experiment; and the reproductive interference parameters $\beta_{TcTu}$ and $\beta_{TuTc}$, the adult offspring sex ratio (SR), total number of sons (Sons), and total number of adult offspring (Offspring) were estimated from the RI experiment.

| | Parameter | | | | | | | | | |
|---|---|---|---|---|---|---|---|---|---|---|
| **Line** | $\lambda_{FC}$ | $\alpha_{TcTc}$ | $\alpha_{TcTu}$ | $\alpha_{TuTc}$ | $\lambda_{RI}$ | $\beta_{TcTu}$ | $\beta_{TuTc}$ | SR | Sons | Offspring |
| 4 | 2.90±0.50 | 0.04±0.02 | 0.04±0.02 | 0.09±0.05 | 4.40±0.61 | 0.28±0.41 | 0.0001 | 0.23±0.06 | 6.1±2.14 | 28.1±7.20 |
| 6 | 5.93±0.89 | 0.17±0.05 | 0.10±0.06 | 0.09±0.04 | 5.54±0.74 | 0.21±0.38 | 0.62±0.63 | 0.28±0.16 | 8.4±1.81 | 36.1±7.23 |
| 8 | 2.40±0.88 | 0.01±0.02 | 0.03±0.03 | 0.11±0.05 | 5.58±0.55 | 0.41±0.49 | 0.14±0.32 | 0.24±0.05 | 8.2±1.48 | 36.1±5.74 |
| 9 | 4.47±0.54 | 0.07±0.03 | 0.07±0.04 | 0.11±0.04 | 4.86±0.28 | 0.56±0.55 | 0.18±0.28 | 0.28±0.08 | 10.5±5.72 | 34.8±5.86 |
| 10 | 0.87±0.34 | 0.0001 | 0.0001 | 0.09±0.05 | 4.00±0.69 | 0.64±0.54 | 0.0001 | 0.24±0.09 | 8±3.47 | 28±9.81 |
| 12 | 3.30±0.63 | 0.08±0.03 | 0.07±0.04 | 0.08±0.03 | 3.98±0.60 | 0.43±0.45 | 0.07±0.44 | 0.20±0.07 | 5.3±2.41 | 25.2±7.52 |
| 14 | 4.93±0.56 | 0.09±0.02 | 0.06±0.04 | 0.16±0.04 | 8.32±0.53 | 0.66±0.63 | 0.20±0.46 | 0.14±0.04 | 6.5±2.09 | 48.1±5.19 |
| 15 | 4.47±0.79 | 0.10±0.03 | 0.09±0.05 | 0.11±0.04 | 8.06±0.58 | 0.67±0.57 | 0.27±0.37 | 0.12±0.03 | 5.8±1.65 | 46.1±6.94 |
| 16 | 3.70±0.84 | 0.07±0.03 | 0.04±0.03 | 0.08±0.04 | 7.08±0.67 | 0.45±0.52 | 0.35±0.64 | 0.15±0.03 | 6±0.83 | 41.4±6.68 |
| 17 | 4.90±1.03 | 0.14±0.05 | 0.16±0.08 | 0.10±0.04 | 5.92±0.56 | 0.28±0.36 | 0.0001 | 0.19±0.05 | 7±2.19 | 36.6±6.25 |
| 18 | 5.27±0.70 | 0.16±0.04 | 0.15±0.08 | 0.11±0.04 | 6.38±0.66 | 0.66±0.52 | 0.0001 | 0.17±0.04 | 6.4±1.66 | 38.3±7.25 |
| 21 | 3.92±0.31 | 0.09±0.03 | 0.08±0.05 | 0.08±0.03 | 6.84±0.56 | 0.39±0.44 | 0.0001 | 0.15±0.03 | 6.1±2.03 | 40.3±7.20 |
| 22 | 3.92±0.98 | 0.07±0.03 | 0.08±0.05 | 0.11±0.04 | 7.10±0.62 | 0.72±0.66 | 0.06±0.17 | 0.15±0.03 | 6.1±1.84 | 41.6±7.19 |
| 25 | 3.48±1.03 | 0.07±0.03 | 0.05±0.04 | 0.10±0.04 | 5.54±0.45 | 0.38±0.46 | 0.17±0.47 | 0.19±0.04 | 6.7±1.96 | 34.4±5.33 |
| 28 | 3.00±0.58 | 0.07±0.03 | 0.06±0.03 | 0.07±0.03 | 4.42±0.76 | 0.46±0.54 | 0.13±0.32 | 0.35±0.19 | 11±5.41 | 33.1±5.18 |
| 29 | 2.92±0.65 | 0.05±0.03 | 0.08±0.04 | 0.12±0.04 | 6.34±1.02 | 0.66±0.59 | 0.0001 | 0.31±0.14 | 11.2±4.51 | 42.9±7.79 |
| 30 | 5.26±0.77 | 0.12±0.03 | 0.18±0.08 | 0.16±0.05 | 3.92±0.63 | 0.43±0.44 | 0.0001 | 0.22±0.05 | 5.2±1.70 | 24.8±7.18 |
| 31 | 4.86±0.45 | 0.10±0.02 | 0.13±0.07 | 0.13±0.05 | 7.72±0.48 | 0.43±0.43 | 0.31±0.48 | 0.16±0.04 | 7.3±2.03 | 45.9±5.27 |
| 32 | 4.67±0.73 | 0.10±0.04 | 0.07±0.05 | 0.09±0.03 | 5.32±0.82 | 0.42±0.48 | 0.05±0.26 | 0.20±0.04 | 6.3±1.89 | 32.9±9.55 |
| 35 | 4.46±0.58 | 0.11±0.05 | 0.07±0.05 | 0.13±0.05 | 7.04±0.57 | 0.86±0.72 | 0.07±0.33 | 0.14±0.03 | 5.8±1.59 | 41±6.76 |
| 36 | 4.42±0.75 | 0.09±0.03 | 0.08±0.05 | 0.15±0.05 | 6.82±0.66 | 0.41±0.44 | 0.04±0.22 | 0.13±0.02 | 5.1±0.94 | 39.2±6.92 |
| 37 | 5.27±0.77 | 0.15±0.05 | 0.17±0.09 | 0.11±0.03 | 6.76±0.46 | 0.40±0.43 | 0.28±0.29 | 0.16±0.03 | 6.7±1.65 | 40.5±5.53 |
| 38 | 5.70±0.70 | 0.21±0.06 | 0.18±0.10 | 0.11±0.03 | 7.52±0.51 | 0.38±0.41 | 0.42±0.60 | 0.14±0.03 | 6.4±1.85 | 44±5.68 |
| 39 | 4.37±0.55 | 0.09±0.02 | 0.06±0.04 | 0.16±0.05 | 5.62±0.46 | 0.36±0.42 | 0.19±0.55 | 0.18±0.07 | 5.9±1.95 | 34±4.25 |
| 40 | 3.13±0.76 | 0.05±0.03 | 0.08±0.04 | 0.11±0.06 | 4.80±0.70 | 0.69±0.58 | 0.0001 | 0.16±0.05 | 4.4±1.58 | 28.4±7.71 |
| 41 | 3.93±0.91 | 0.11±0.04 | 0.08±0.05 | 0.11±0.05 | 7.92±0.88 | 0.36±0.41 | 0.13±0.32 | 0.22±0.13 | 9.4±4.09 | 49±5.28 |
| 42 | 5.42±0.57 | 0.20±0.05 | 0.19±0.11 | 0.10±0.04 | 7.98±0.41 | 0.54±0.52 | 1.00±1.00 | 0.17±0.05 | 8.3±3.17 | 48.2±4.49 |
| 43 | 3.70±0.78 | 0.07±0.03 | 0.07±0.04 | 0.15±0.04 | 4.46±0.70 | 0.0001 | 0.0001 | 0.32±0.16 | 10.2±4.60 | 32.5±7.17 |
| 44 | 3.27±0.68 | 0.05±0.02 | 0.06±0.03 | 0.13±0.04 | 5.34±0.39 | 0.24±0.38 | 0.27±0.40 | 0.24±0.08 | 9±3.90 | 35.7±5.81 |

**Figure SI1. Estimated parameter values from the offspring production data of the FC experiment.** Dots represent the mean parameter value of each Tc line for (A) the intrinsic growth rate, λ; (B) the sensitivity of Tc to conspecific competitors, $\alpha_{TcTc}$; (C) the sensitivity of Tc to heterospecific competitors, $\alpha_{TcTu}$; and (D) the effect of Tc on heterospecific competitors, $\alpha_{TuTc}$. Lines are sorted by increasing value for each estimated parameter, such that their order differs among figures. Error bars represent the 95% confidence interval.

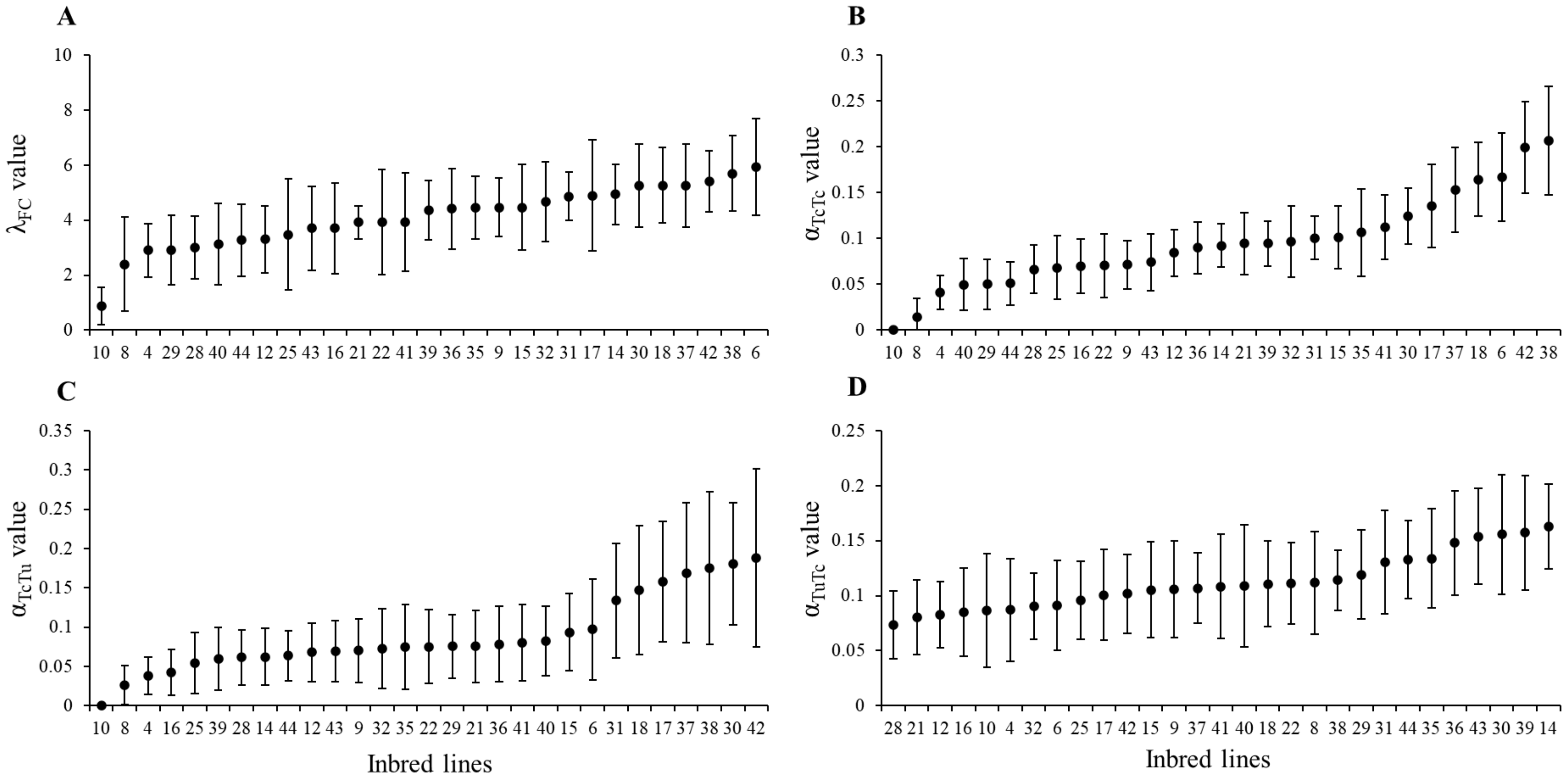

**Figure SI2. Estimated parameters and variables from the offspring production data of the RI experiment.** Dots represent the mean value of each Tc line for (A) the intrinsic growth rate, $\lambda$; (B) the adult offspring sex ratio (computed as the proportion of sons in adult offspring); (C) the total number of adult sons; (D) total number of adult offspring (computed as the sum of adult sons and daughters); (E) the sensitivity of Tc to reproductive interference by Tu, $\beta_{TcTu}$; and (F) the sensitivity of Tu to reproductive interference by Tc, $\beta_{TuTc}$. Lines are sorted by increasing value for each estimated parameter, such that their order differs among figures. Error bars represent the 95% confidence interval.

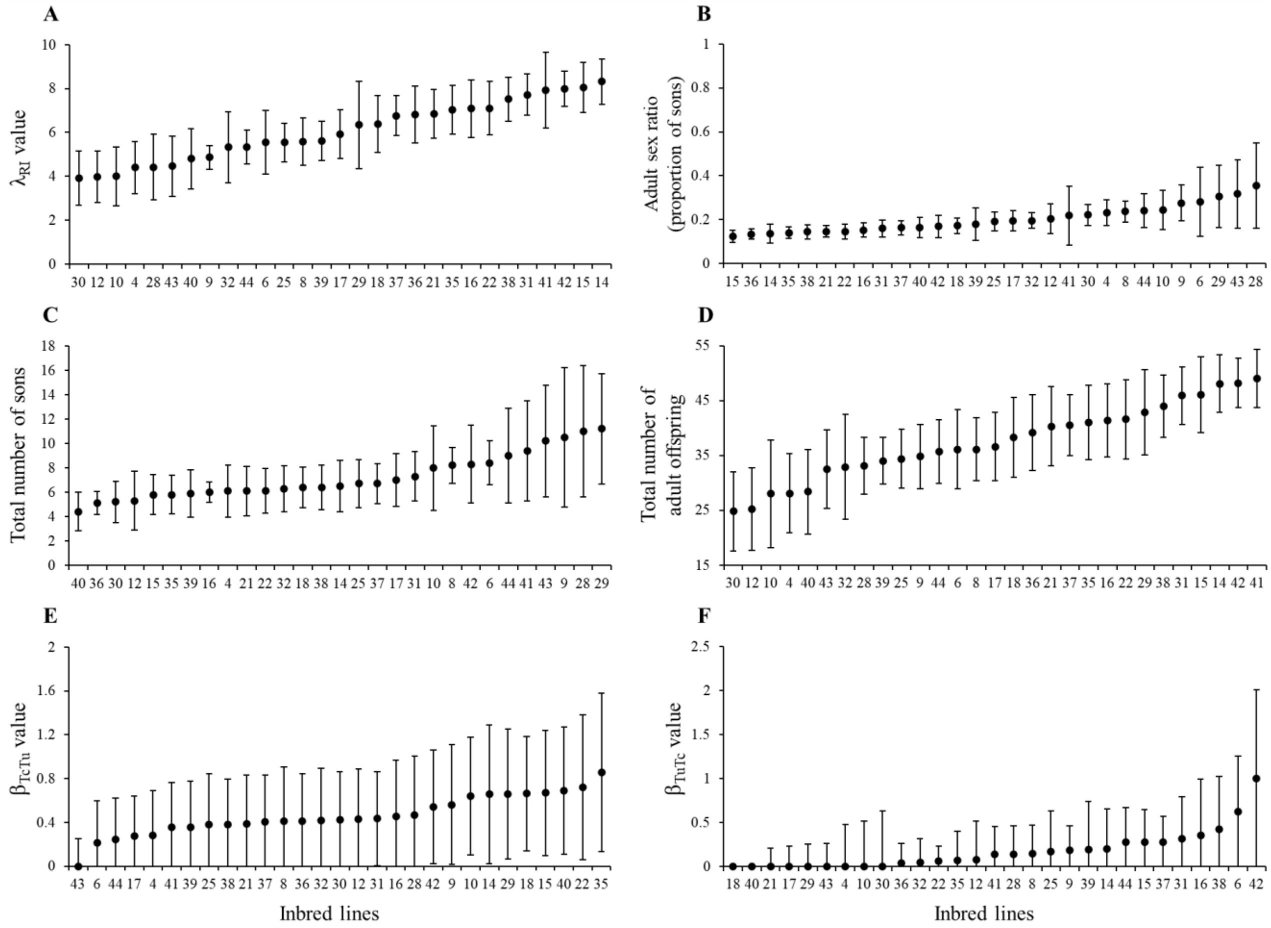

**Table SI2. Estimates of heritability for each parameter or variable in the FC and RI experiments.** Broad-sense heritability ($H^2$) was estimated from the offspring production data, by sub-setting data obtained for each combination of focal individuals and competitors (FC experiment) or heterospecific mates (RI experiment; see Methods for a detailed explanation). The 95% highest posterior density interval (HPDI) and the difference between the deviance information criteria (ΔDIC) of models with or without 'line' as random factor are also provided for each estimate. ΔDIC values above 2 (shaded in grey) indicate that the model accounting for 'line' has a significantly better fit (*i.e.*, the assessed parameter or variable is heritable). SR: Adult offspring sex ratio, computed as the proportion of sons. Sons: Total number of adult sons. Offspring: Total number of adult sons and daughters.

| Data subset | Parameter/variable | $H^2$ | HPDI | ΔDIC |
|---|---|---|---|---|
| Tc females alone | $\lambda_{FC}$ | 0.13 | 0.03, 0.24 | 20.79 |
| | $\lambda_{RI}$ | 0.20 | 0.08, 0.33 | 44.47 |
| Tc females exposed to conspecifics | $\alpha_{TcTc}$ | 0.07 | 0.02, 0.11 | 49.57 |
| Tc females exposed to heterospecifics | $\alpha_{TcTu}$ | 0.06 | 0.02, 0.11 | 36.63 |
| Tu females exposed to heterospecifics | $\alpha_{TuTc}$ | 0.01 | 0.001, 0.03 | -0.38 |
| Tc females exposed to heterospecifics | $\beta_{TcTu}$ | 0.06 | 0.0007, 0.15 | 7.22 |
| Tu females exposed to heterospecifics | $\beta_{TuTc}$ | 0.05 | 0.0006, 0.12 | 3.91 |
| Tc females alone | SR | 0.14 | 0.04, 0.23 | 31.56 |
| | Sons | 0.05 | 0.001, 0.12 | 4.56 |
| | Offspring | 0.16 | 0.06, 0.28 | 22.74 |

**Table SI3. Correlations between estimated parameters and variables.** Correlations were tested among all possible pairs of parameters for which heritability was significant (*i.e.*, correlations with $\alpha_{TuTc}$ were not assessed, cf. Table SI1). FDR corrections were used to account for multiple testing. Rows shaded in grey display significant correlations at the 5% level. *r*: Pearson correlation coefficient. SR: Adult offspring sex ratio, computed as the proportion of sons. Sons: Total number of adult sons. Offspring: Total number of adult sons and daughters.

| **Parameter pairs** | ***r*** | **p-value (FDR)** |
|---|---|---|
| $\alpha_{TcTc}$ and $\alpha_{TcTu}$ | 0.86 | <0.0001 |
| $\alpha_{TcTc}$ and $\lambda_{FC}$ | 0.89 | <0.0001 |
| $\alpha_{TcTu}$ and $\lambda_{FC}$ | 0.78 | <0.0001 |
| $\beta_{TcTu}$ and $\beta_{TuTc}$ | -0.22 | 0.21 |
| $\beta_{TcTu}$ and $\lambda_{RI}$ | 0.31 | 0.08 |
| $\beta_{TuTc}$ and $\lambda_{RI}$ | 0.35 | 0.05 |
| $\alpha_{TcTc}$ and $\beta_{TcTu}$ | -0.09 | 0.38 |
| $\alpha_{TcTu}$ and $\beta_{TcTu}$ | -0.04 | 0.41 |
| $\alpha_{TcTc}$ and $\beta_{TuTc}$ | 0.42 | 0.04 |
| $\alpha_{TcTu}$ and $\beta_{TuTc}$ | 0.17 | 0.26 |
| $SR_{RI}$ and $\beta_{TcTu}$ | -0.38 | 0.03 |
| $SR_{RI}$ and $\beta_{TuTc}$ | -0.07 | 0.39 |
| Offspring and $\lambda_{RI}$ | 0.96 | <0.0001 |
| $SR_{RI}$ and Offspring | -0.42 | 0.03 |
| Offspring and Sons | 0.15 | 0.29 |
| $SR_{RI}$ and $\lambda_{RI}$ | -0.64 | <0.0001 |
| $\lambda_{FC}$ and $\lambda_{RI}$ | 0.48 | 0.02s |

**Figure SI3. Correlations among offspring production parameters and variables.** Panels display the correlations between (A) intrinsic growth and total number of adult offspring (sum of adult sons and daughters); (B) number of adult sons and total number of adult offspring; and (C) total number of adult offspring and the offspring sex ratio (proportion of males in adult offspring) obtained in the RI experiment. Panel (D) shows correlations between intrinsic growth rates measured in the two experiments. Each dot represents the mean of the values measured for a single Tc line. Error bars represent the 95% confidence interval. Red trendlines indicate a significant correlation between parameters at the 5% level.

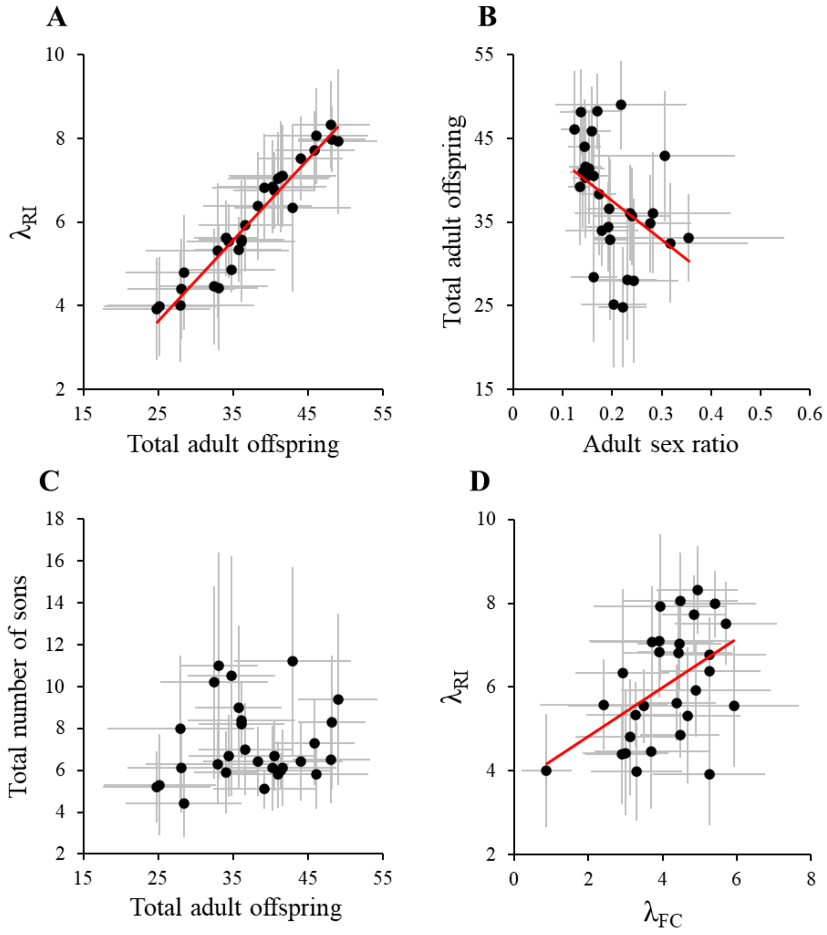
