## Supplementary materials II for "Spider mite genotypes with higher growth rate suffer more from competition but exert stronger reproductive interference"

### Measuring adult female and male body size

To test whether adult body size could play a role in determining the negative impact of competition for food and reproductive interference on spider mite population growth, we measured the size of individual mites, both males and females, from a set of 18 out of the 29 Tc lines, and tested for genetic correlations with variables involved in interspecific interactions or offspring production. The procedures to do so and results obtained are described below.

### Materials and Methods

Measurements were conducted independently for females and males. The protocol was identical for both sexes (described below), differing only in the age cohort procedure used to obtain enough individuals of the same age that developed in the exact same environmental conditions.

*Female size cohorts*

Eleven days prior to the measurements, age cohorts were performed to obtain the adult females used for body size measurements: adult females taken from the population cages with expanded lines were installed in groups of 25 per line on large leaf fragments placed on water-soaked cotton Petri dishes. Two days prior to the measurements, at least 5 quiescent females from the age cohorts of each line were isolated on small leaf fragments until emerging as adults, to ensure they remained virgin and were of similar age. As all 18 Tc lines could not be tested simultaneously, a total of 6 experimental blocks were carried out, with odd-numbered blocks (1, 3 and 5) containing 2 to 9 replicates of 10 of the lines and even-numbered blocks (2, 4 and 6) containing similar numbers of replicates of the remaining 8 lines. This resulted in a minimum of 7 and a maximum of 20 experimental replicates per line, for a total of 299 replicates.

*Male size cohorts*

Ten days prior to the measurements, age cohorts were performed to obtain the adult males used for body size measurements. These consisted of 5 quiescent females of each line, installed on large leaf fragments placed on water-soaked cotton in separate Petri dishes. 2 days prior to the measurements, at least 5 quiescent males from the age cohorts of each line were isolated on small leaf fragments until emerging as adults, to ensure they were of similar age. As all 18 Tc lines could not be tested simultaneously, a total of 7 experimental blocks were carried out. Due to issues of low population growth for several lines, between 8 and 13 different lines were tested in each block, containing 1 to 10 replicates of each line. This resulted in a minimum of 3 and a maximum of 16 experimental replicates per line, for a total of 232 replicates.

*Procedure for measuring spider mite body size*

Individual spider mites were placed on a small droplet of water in a transparent Petri dish. The dish was then placed under a Leica S9E stereomicroscope over a white background, and a photograph was captured using a Leica MTU-293 camera adapter and an ImagingSource® DFK 23UX174 colour camera. A micrometer calibration slide was photographed at the start of each session for scale. The photographs were later used to measure the length and width of the idiosome of each individual mite (as exemplified in Figure SII1; Enders 1993) using the software ImageJ (Clotuche et al. 2012). Each individual was measured 3 times independently to account for observation errors, and the mean of the 3 measurements was used for analyses.

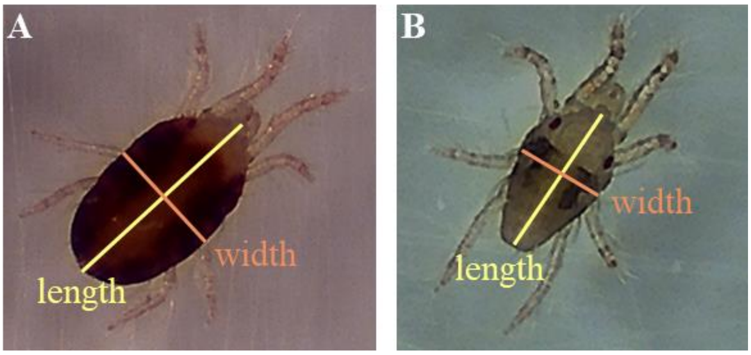


**Figure SII1. Example of body size measurements of female and male spider mites.** Lines display the length and width of the idiosome, in yellow and orange, respectively, measured on photographs of an adult female (A) and an adult male (B) spider mite.

*Statistical analyses*

Analyses were carried out using the R statistical software (v4.2.2), following the same procedures as those described in the main manuscript. Body area for adult females and males was calculated as the area of an ellipse based on the idiosome length and width (*i.e.*, as [(idiosome length/2) * (idiosome width/2) * π]; Table SII1; Figure SII2). Broad-sense heritability ($H^2$) was estimated as described in the main manuscript, for body length, width and area, using their measurements in millimetres (following a log-transformation) as response variables. Correlations were initially tested among different body size measurements for each sex (*e.g.*, female length and width, male width and area, and so on). As these variables were found to be strongly correlated (see below), only the body area of females and males was used to test for genetic correlations (see Figure 1, Table SI2).

**Table SII1. Adult female and male body size for each of the 18 measured Tc lines.** The mean (± 95% confidence interval) body length, width, and area of adult females (♀) and males (♂), in millimetres, measured in the body size experiments are provided for each Tc line.

| | **Parameter** | | | | | |
|---|---|---|---|---|---|---|
| **Line** | ♀ length | ♀ width | ♀ area | ♂ length | ♂ width | ♂ area |
| 4 | 0.435±0.017 | 0.271±0.007 | 0.093±0.006 | 0.276±0.006 | 0.160±0.003 | 0.035±0.001 |
| 6 | 0.433±0.009 | 0.264±0.007 | 0.090±0.004 | 0.261±0.008 | 0.150±0.003 | 0.031±0.001 |
| 8 | 0.418±0.011 | 0.260±0.006 | 0.086±0.004 | 0.256±0.010 | 0.147±0.003 | 0.030±0.002 |
| 10 | 0.420±0.013 | 0.263±0.005 | 0.087±0.004 | 0.277±0.005 | 0.166±0.015 | 0.036±0.003 |
| 12 | 0.434±0.012 | 0.275±0.006 | 0.094±0.005 | 0.267±0.004 | 0.158±0.003 | 0.033±0.001 |
| 14 | 0.437±0.014 | 0.279±0.008 | 0.096±0.006 | 0.265±0.007 | 0.156±0.002 | 0.032±0.001 |
| 15 | 0.444±0.011 | 0.285±0.007 | 0.100±0.005 | 0.276±0.007 | 0.158±0.002 | 0.034±0.001 |
| 28 | 0.433±0.010 | 0.275±0.005 | 0.094±0.003 | 0.258±0.008 | 0.157±0.003 | 0.032±0.001 |
| 29 | 0.415±0.008 | 0.268±0.004 | 0.087±0.003 | 0.269±0.006 | 0.156±0.002 | 0.033±0.001 |
| 30 | 0.422±0.018 | 0.268±0.007 | 0.089±0.006 | 0.264±0.005 | 0.157±0.002 | 0.032±0.001 |
| 32 | 0.404±0.010 | 0.262±0.007 | 0.084±0.004 | 0.260±0.003 | 0.153±0.003 | 0.031±0.001 |
| 35 | 0.381±0.015 | 0.245±0.008 | 0.074±0.005 | 0.262±0.006 | 0.152±0.002 | 0.031±0.001 |
| 37 | 0.406±0.011 | 0.263±0.004 | 0.084±0.003 | 0.272±0.003 | 0.156±0.004 | 0.033±0.001 |
| 38 | 0.401±0.006 | 0.257±0.005 | 0.081±0.002 | 0.265±0.007 | 0.154±0.003 | 0.032±0.001 |
| 40 | 0.415±0.014 | 0.262±0.005 | 0.086±0.004 | 0.262±0.005 | 0.153±0.002 | 0.031±0.001 |
| 42 | 0.432±0.012 | 0.272±0.007 | 0.092±0.005 | 0.265±0.005 | 0.155±0.002 | 0.032±0.001 |
| 43 | 0.399±0.028 | 0.262±0.007 | 0.082±0.008 | 0.257±0.007 | 0.149±0.002 | 0.030±0.001 |
| 44 | 0.414±0.010 | 0.262±0.004 | 0.085±0.003 | 0.265±0.007 | 0.149±0.003 | 0.031±0.002 |

**Figure SII2. Adult female and male body size for each of the 18 measured Tc lines.** Dots represent the mean female length (A), width (B), and body area (C), as well as the mean male length (D), width (E), and body area (F) of each Tc line. Error bars represent the 95% confidence interval.

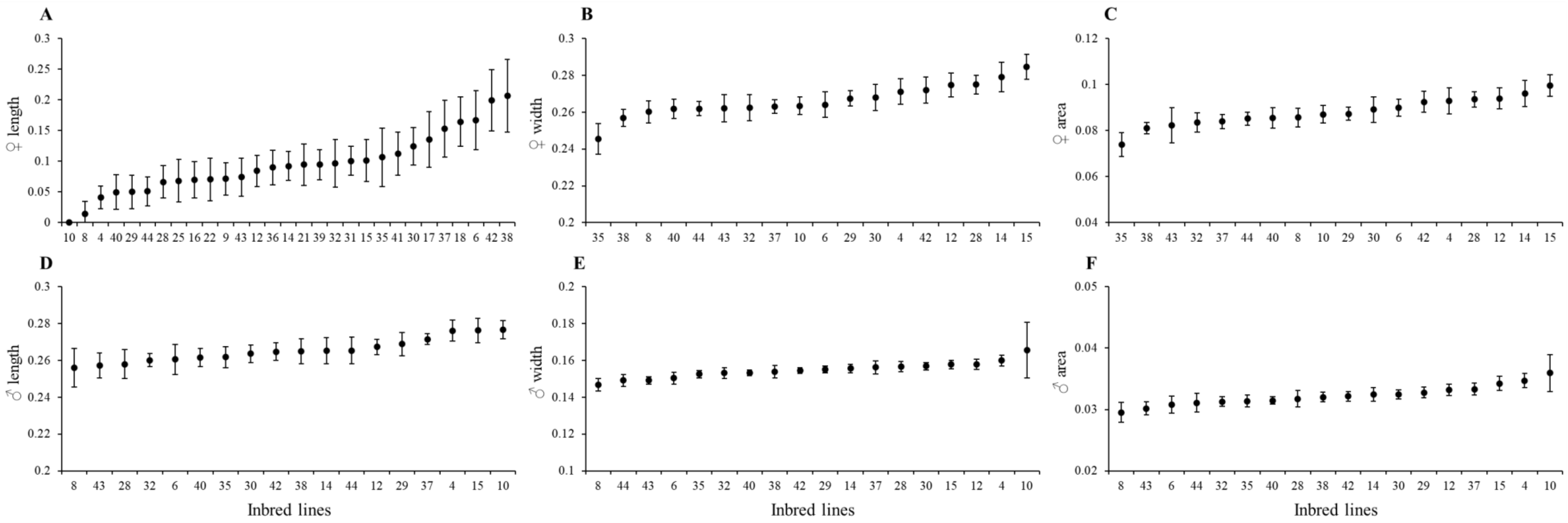

**Results**

*Broad-sense heritability*

We found significant broad-sense heritability for all body size variables, except male body area (Table SII2; Figure SII3).

**Table SII2. Estimates of heritability for adult female and male body size.** The estimate of broad-sense heritability ($H^2$), the 95% highest posterior density interval (HPDI) and the difference between the deviance information criteria (ΔDIC) of models with or without 'line' as random factor are provided for each body size measurement. ΔDIC values above 2 (shaded in grey) indicate that the model accounting for 'line' was significantly different from the model excluding it (*i.e.*, the assessed variable is heritable).

| Variable | $H^2$ | HPDI | ΔDIC |
|---|---|---|---|
| ♀ length | 0.204 | 0.042, 0.394 | 37.54 |
| ♀ width | 0.230 | 0.040, 0.443 | 55.87 |
| ♀ area | 0.230 | 0.032, 0.438 | 47.38 |
| ♂ length | 0.250 | 0.062, 0.457 | 15.97 |
| ♂ width | 0.249 | 0.048, 0.462 | 8.21 |
| ♂ area | 0.254 | 0.043, 0.481 | -3.02 |

**Figure SII3. Estimates of heritability for adult female and male body size.** Dots represent mean broad-sense heritability for each body size measurement. Error bars represent the 95% highest posterior density interval (HPDI). Asterisks indicate values of heritability significantly different from 0 at the 5% level.

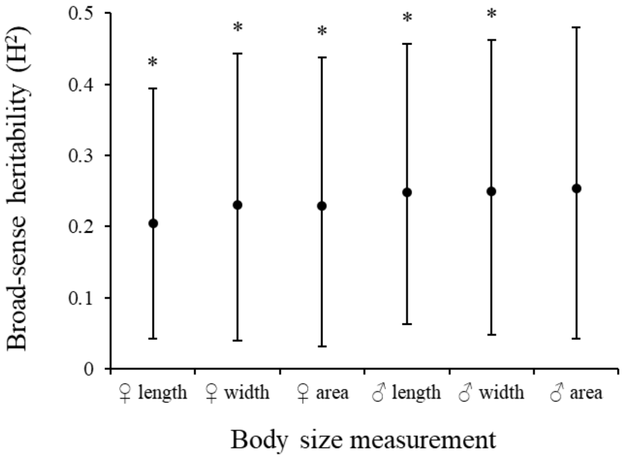


*Genetic correlations*

We found significant highly positive correlations among the different female size measurements (♀ length and ♀ width, $r = 0.90$, $P < 0.0001$; ♀ length and ♀ area, $r = 0.98$, $P < 0.0001$; ♀ width and ♀ area, $r = 0.97$, $P < 0.0001$) and among male length and width, ($r = 0.78$, $P < 0.0001$). However, female body area and male length were not genetically correlated (♀ area and ♂ length, $r = 0.37$, $P = 0.16$; Table SII3; Figure SII4). Furthermore, we found no significant correlations between body area of females and male length and any parameter related to offspring production (intrinsic growth, λ, in either the FC or the RI experiments, total offspring production, or offspring sex ratio), to food competition (intraspecific competition, $\alpha_{TcTc}$, and interspecific competition, $\alpha_{TcTu}$), or to reproductive interference (the ability of Tc to cope with heterospecific mates, $\beta_{TcTu}$, or the effect of Tc on heterospecific mates, $\beta_{TcTu}$; Table SII3; see also Figures SII5 and SII6).

**Table SII3. Correlations among body size and interaction parameters estimated or offspring production variables obtained from the FC and RI experiments.** Correlations were tested among female and male body size measurements, as well as between female body area and male length and all other parameters for which heritability was significant (*i.e.*, correlations with $\alpha_{TuTc}$ were not assessed). FDR corrections were used to account for multiple testing. Rows shaded in grey display significant correlations at the 5% level. *r*: Pearson correlation coefficient. SR: Offspring sex ratio, computed as the proportion of sons. Total offspring: sum of adult sons and daughters.

| **Variable pairs** | ***r*** | **p-value (FDR)** |
|---|---|---|
| ♀ length and ♀ width | 0.90 | <0.0001 |
| ♀ length and ♀ area | 0.98 | <0.0001 |
| ♀ width and ♀ area | 0.97 | <0.0001 |
| ♂ length and ♂ width | 0.78 | <0.0001 |
| ♀ area and ♂ length | 0.37 | 0.30 |
| ♀ area and $\lambda_{FC}$ | -0.03 | 0.50 |
| ♀ area and $\lambda_{RI}$ | 0.004 | 0.50 |
| ♀ area and total offspring | -0.01 | 0.50 |
| ♀ area and SR | 0.03 | 0.50 |
| ♀ area and $\alpha_{TcTc}$ | -0.07 | 0.50 |
| ♀ area and $\alpha_{TcTu}$ | -0.05 | 0.50 |
| ♀ area and $\beta_{TcTu}$ | 0.05 | 0.50 |
| ♀ area and $\beta_{TuTc}$ | 0.09 | 0.50 |
| ♂ length and $\lambda_{FC}$ | -0.20 | 0.50 |
| ♂ length and $\lambda_{RI}$ | 0.12 | 0.50 |
| ♂ length and total offspring | 0.06 | 0.50 |
| ♂ length and SR | -0.31 | 0.35 |
| ♂ length and $\alpha_{TcTc}$ | -0.10 | 0.50 |
| ♂ length and $\alpha_{TcTu}$ | -0.05 | 0.50 |
| ♂ length and $\beta_{TcTu}$ | 0.27 | 0.40 |
| ♂ length and $\beta_{TuTc}$ | -0.07 | 0.50 |

**Figure SII4. Correlations among body size measurements.** Panels display the correlations between female body length and width (A), length and area (B), and width and area (C), between male body length and width (D), and between female body area and male length (G). Each dot represents the mean of the values obtained for a single Tc line. Error bars represent the 95% confidence interval. Red trendlines indicate a significant correlation between variables at the 5% level.

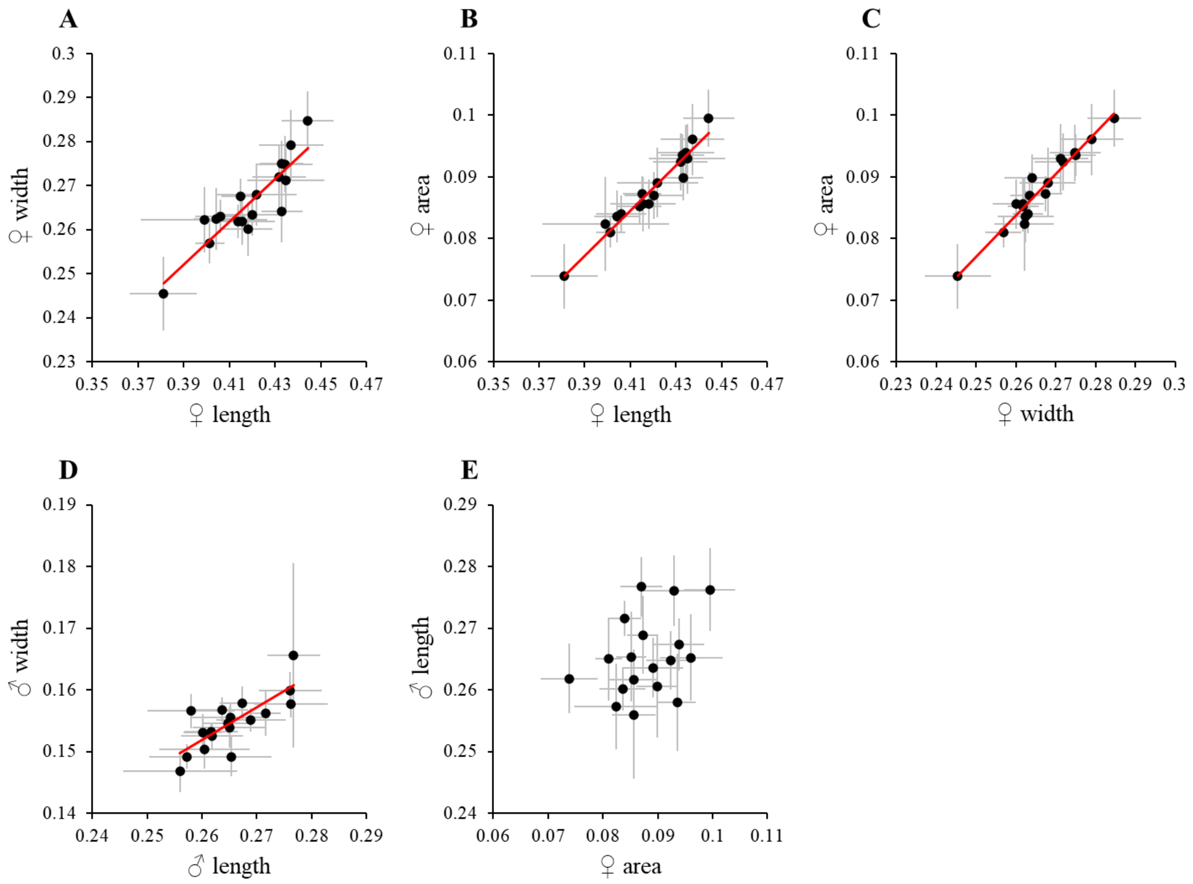

**Figure SII5. Correlations among body area and offspring production variables.** The top row of panels displays the correlations between female body area and: intrinsic growth in either the FC (A) or the RI (B) experiments, total adult offspring production (C), and adult offspring sex ratio, *i.e.* proportion of sons (D). The bottom row of panels (E through H) displays the same correlations for male length. Each dot represents the mean of the values obtained for a single Tc line. Error bars represent the 95% confidence interval.

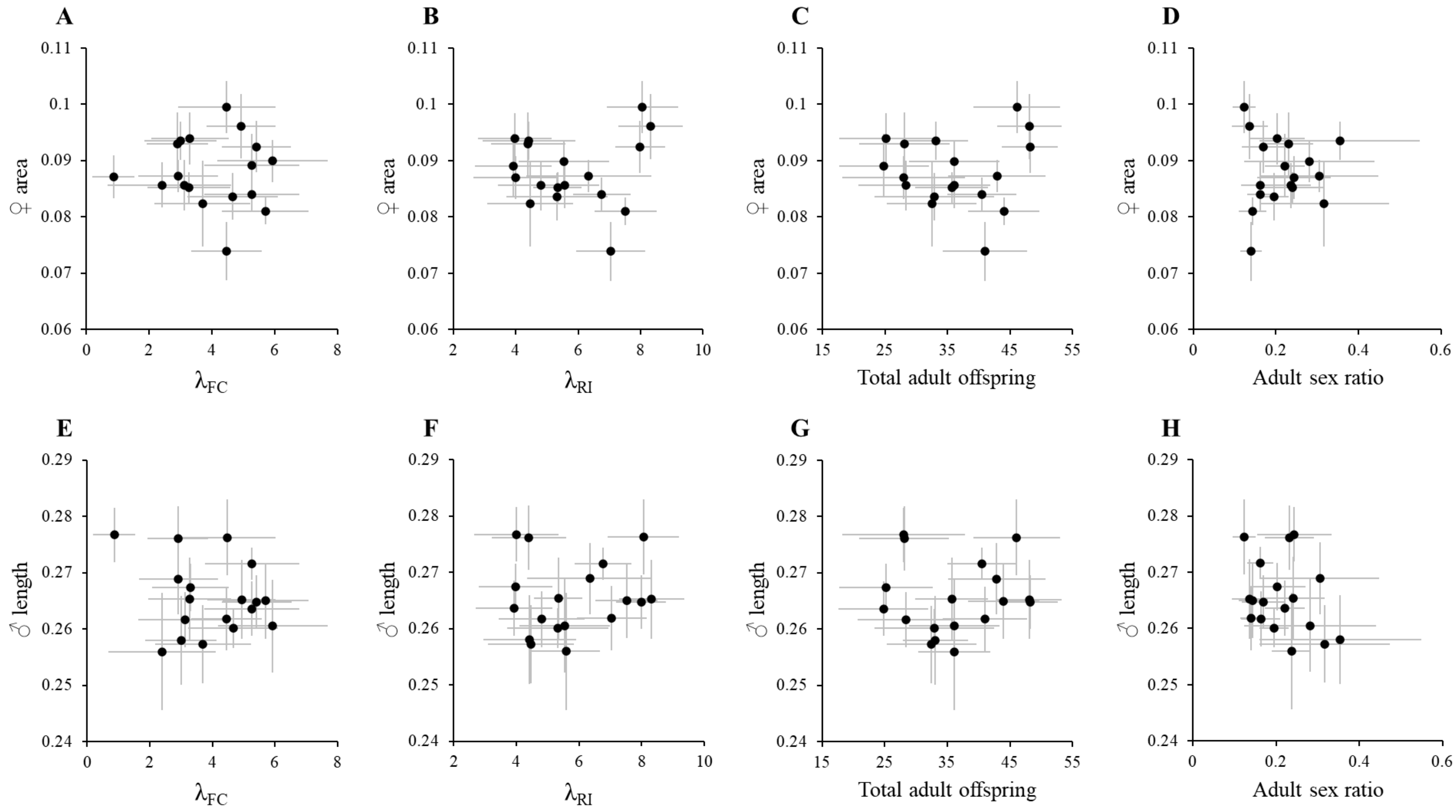

**Figure SII6. Correlations among body area and interspecific interaction parameters.** The top row of panels displays the correlations between female body area and sensitivity to conspecific (A) or heterospecific (B) competitors ($\alpha_{TcTc}$ and $\alpha_{TcTu}$ respectively), sensitivity to heterospecific mates, $\beta_{TcTu}$ (C), and the effect of Tc on heterospecific mates, $\beta_{TuTc}$ (D). The bottom row of panels (E through H) displays the same correlations for male length. Each dot represents the mean of the values obtained for a single Tc line. Error bars represent the 95% confidence interval.

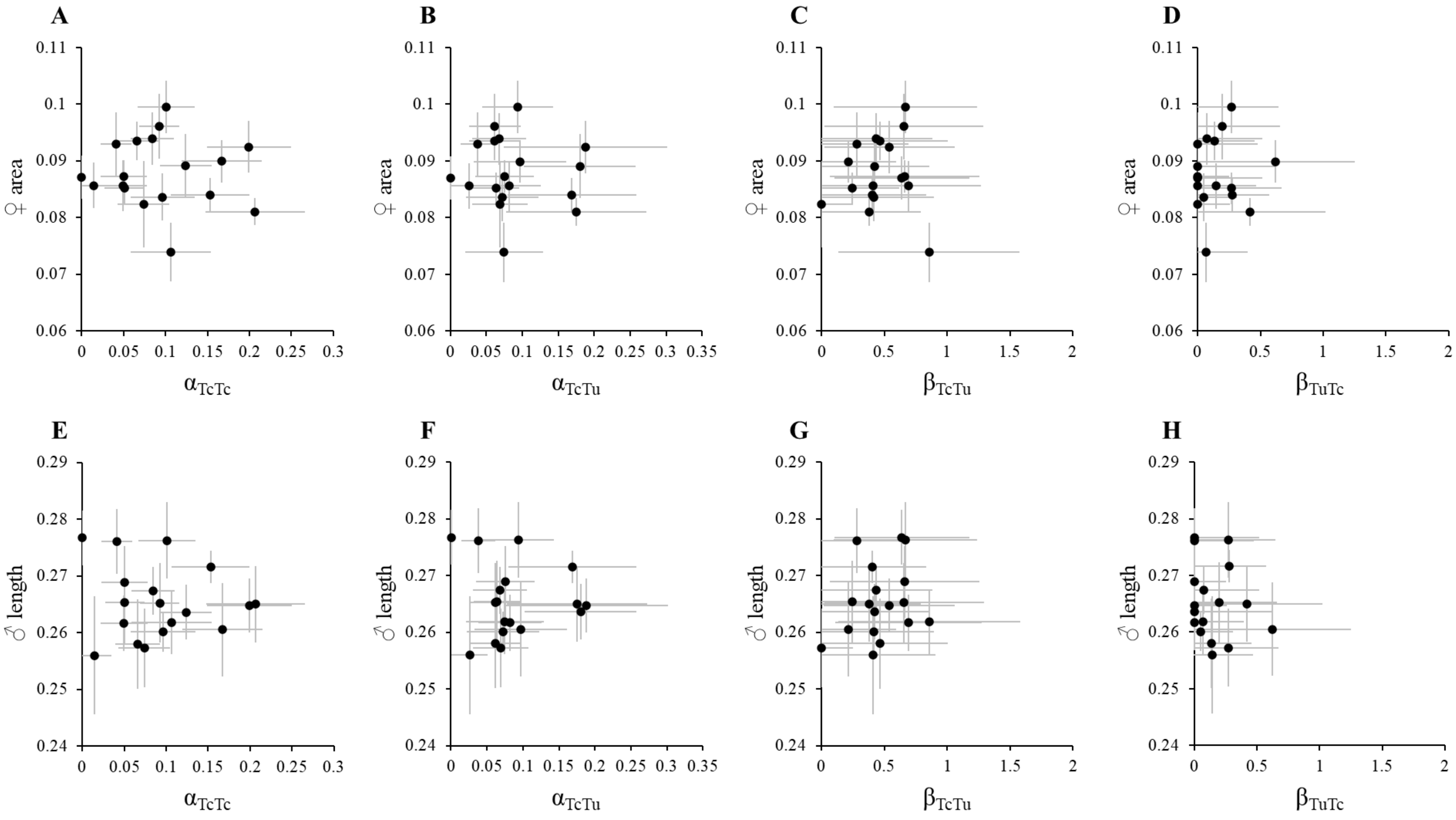

References

Clotuche, G., A.-C. Mailleux, J.-L. Deneubourg, C. Detrain, and T. Hance. 2012. The silk road of *Tetranychus urticae*: is it a single or a double lane? Exp. Appl. Acarol. 56:345–354.

Enders, M. M. 1993. The effect of male size and operational sex ratio on male mating success in the common spider mite, *Tetranychus urticae* Koch (Acari: Tetranychidae). Anim. Behav. 46:835–846.
